# Interpretable machine learning coupled to gene regulatory networks uncovers subcircuits underlying cell fate decisions

**DOI:** 10.64898/2026.09.28.754797

**Authors:** Zarifeh H Rarani, Swapnil Keshari, Akanksha Sachan, Ankur Saini, Nicholas Pease, Jingyu Fan, Alayna Fu, Yuchen C Liu, Aditi U Gurkar, Greg M Delgoffe, Harinder Singh, Jishnu Das

## Abstract

Gene regulatory networks (GRNs) model causal linkages that control cell fate decisions and differentiation transitions. Prioritizing regulatory subnetworks underlying cell state differences is of critical importance, but current methods including those reliant on topological metrics introduce circularity as the metrics prioritizing TFs are computed from the same networks whose assumptions they inherit. Separately, interpretable machine learning methods can identify latent factors (LFs) that discriminate cellular states with formal statistical guarantees but do not model regulatory linkages. Here, we present FOCAL (Factor-Outcome Coupling for Assessment of Linkages), a paradigm to prioritize regulatory subnetworks by coupling state-specific and dynamic GRNs with outcome-supervised LFs learned using interpretable machine learning without reference to network topology. This shifts GRN focus from macroscopic TF nodes to state-specific and dynamic TF-gene linkages. In B and T cells, FOCAL identified GIFs (GRNs coupled to Interpretable latent Factors), prioritized regulatory subnetworks underlying established states as well as transient regulatory episodes preceding them. By coupling LFs learnt from perturbation experiments of lineage-defining TFs, FOCAL identified transcriptional predisposition to alternative fates within progenitor cell populations before overt differentiation. This uncovered a novel NFATC2-IRF8 interplay in activated B cells, that was validated by in-vitro and in-vivo genetic perturbations. The two transcription factors act cooperatively to restrain extrafollicular plasmablast differentiation and promote germinal center B cell fate.

## Introduction

Cellular developmental and differentiation transitions are orchestrated by gene regulatory networks (GRNs) comprising lineage-determining and function-controlling transcription factors (TFs) and their target genes. State-specific and dynamic GRNs are being widely assembled from multi-omic datasets by combining covariation in transcript abundance and chromatin accessibility with TF motif enrichment or binding information^1^. The sequence-directed integration step is crucial, as it assigns candidate regulatory directions to the statistical associations and configures them in sequence- and accessibility-constrained models, as TF to cis-regulatory element (CRE) to gene linkages. In doing so, the overall framework enables the construction of densely linked regulatory graphs that can generate testable hypotheses about the control of gene activity and cell-fate dynamics. Such models can be used to simulate perturbations of CREs or TFs and predict their effects on individual genes or genomic programs (MIRA^2^, CellOracle^3^, Dictys^4^). Once cell-state annotations are applied, these simulations also provide criteria for ranking TFs by their inferred influence on fate transitions^3^. These criteria include the centrality of a TF within a state-specific network, the aggregate effect of in silico deletion of CREs linked to a TF on cell state-associated genes, and the impact of propagating a TF perturbation through the GRN.

The comprehensiveness and complexity of these GRN models necessitate the prioritization of regulatory nodes and edges which however are associated with considerable uncertainties including peak-to-gene assignments of accessible chromatin regions and covariance-based edge inferences. Furthermore, the sequence-anchored layer that provides candidate molecular mechanisms is an additional source of uncertainty. Simple and redundant TF motifs occur frequently and are weakly predictive of genome occupancy and transcriptional regulation. Such motifs delineate TF families rather than individual TFs and therefore distinguishing the actions of paralogs or isoforms, the latter generated by alternative splicing is challenging. These limitations imply that a GRN containing >10□ edges may include a substantial fraction of linkages that are incorrect, indirect, or inconsequential. Critically, TF prioritization criteria such as centrality^3^ or regulon scores^5,6^ generate circularity in reasoning as these metrics cannot independently detect such errors, because each is calculated from the network itself. By quantifying a TF’s influence based on the coordinated expression of targets already assigned to it by the network, the evaluation metric becomes conditional on the assumed topology. It inherits the sequence and covariance assumptions used to construct the GRN rather than independently testing them. An edge that is misassigned but highly weighted may consequently be scored as influential by each of these measures. Importantly, topological circularity does not arise during this initial GRN assembly process, which serves as a necessary hypothesis-generation step. Rather, it is introduced during the prioritization of nodes and edges. Thus, current GRN assembly frameworks lack a means of assessing the regulatory relevance of nodes and edges using evidence generated independently of the network topology and the sequence priors used to construct them.

Two lines of work have attempted to address this problem by different means. The first utilizes dynamical modeling frameworks such as neural ordinary differential equations^7^, RNA-velocity vector fields^8^, or perturbation simulations (Dynamo^9^, scVelo^8^) to reconstruct continuous state transitions. While these approaches can successfully estimate context-dependent TF activity and identify candidate drivers of developmental trajectories, they largely operate at the level of the macroscopic cell-state manifold. Consequently, their cell-fate predictions are focused on TFs but not fully resolved at the level of individual, testable regulatory linkages, leaving many underlying GRN connections unevaluated.

The second set of approaches uses phenotype-associated gene signatures to prioritize regulators within an inferred network. Master regulator analysis tests whether the targets of a TF are enriched within a signature that distinguishes cellular states (MARINa^10^), and TF activity inference generalizes this by estimating regulator activity from the coordinated expression of its targets rather than from its own transcript (VIPER^11^, DoRothEA^12^/decoupleR^13^). Regulon activity scoring applies a related logic at single-cell resolution (AUCell/SCENIC^5^). However, while these methods incorporate phenotypic contrast, the TF prioritizing signal relies on the expression of targets already assigned to it within the *in-silico* network. Because this inference remains conditional on the assumed network topology, these methods suffer from the circularity described above. They cannot independently test the underlying linkages based on evidence outside of the priors used to build them.

Nevertheless, these complementary approaches establish two principles that motivated our framework: (i) TF activity can be more effectively inferred from the coordinated behavior of its target genes than from its own transcript abundance, and (ii) outcome-associated transcriptional programs can be used to prioritize regulatory components within a large network. To provide an evidence layer based on outcome supervision that assesses the regulatory relevance of edges independently of network topology, we use an interpretable ML model, SLIDE, previously developed by us^14^. It identifies standalone and interacting latent factors (LFs) associated with a cellular state or phenotypic outcome, with formal guarantees concerning LF identifiability, statistical inference, and false-discovery-rate control under the assumptions of the model (Fig. 1). Our approach prioritizes regulatory subcircuits by intersecting GRNs with outcome-supervised latent factors without reference to network topology. The GRNs define candidate causal regulatory relationships, whereas the latent factors identify the transcriptional programs that carry information about the phenotypic contrast. Their intersection therefore prioritizes regulatory gene programs and candidate linkages supported by both mechanistic and outcome-associated evidence. We note that GRN inference frameworks and SLIDE differ along two distinct axes. First, they rest on different forms of evidence. GRN edges carry sequence, accessibility, or TF binding support, whereas SLIDE-selected LFs are delineated using cell-state labels. Second, they differ in how the cellular state contrast is used. Cell-state annotations enter GRN analysis by partitioning cells before or after network construction, and may be used subsequently to score edges, but the contrast between alternative states does not determine the assignment or direction of individual TF to CRE to target gene edges. In distinction, SLIDE selects LFs based on variation that carries information about the cellular states. Though the two analytical frameworks, in part operate on the same transcriptomic data, their independence lies in the nature and assumptions of the modeling approaches. We note that the formal guarantees described above apply to the selection of LFs. The enrichment of TF target sets within state-specific GRNs constitutes a separate inferential step, for which multiple-testing controls are applied independently.

**Figure 1:**
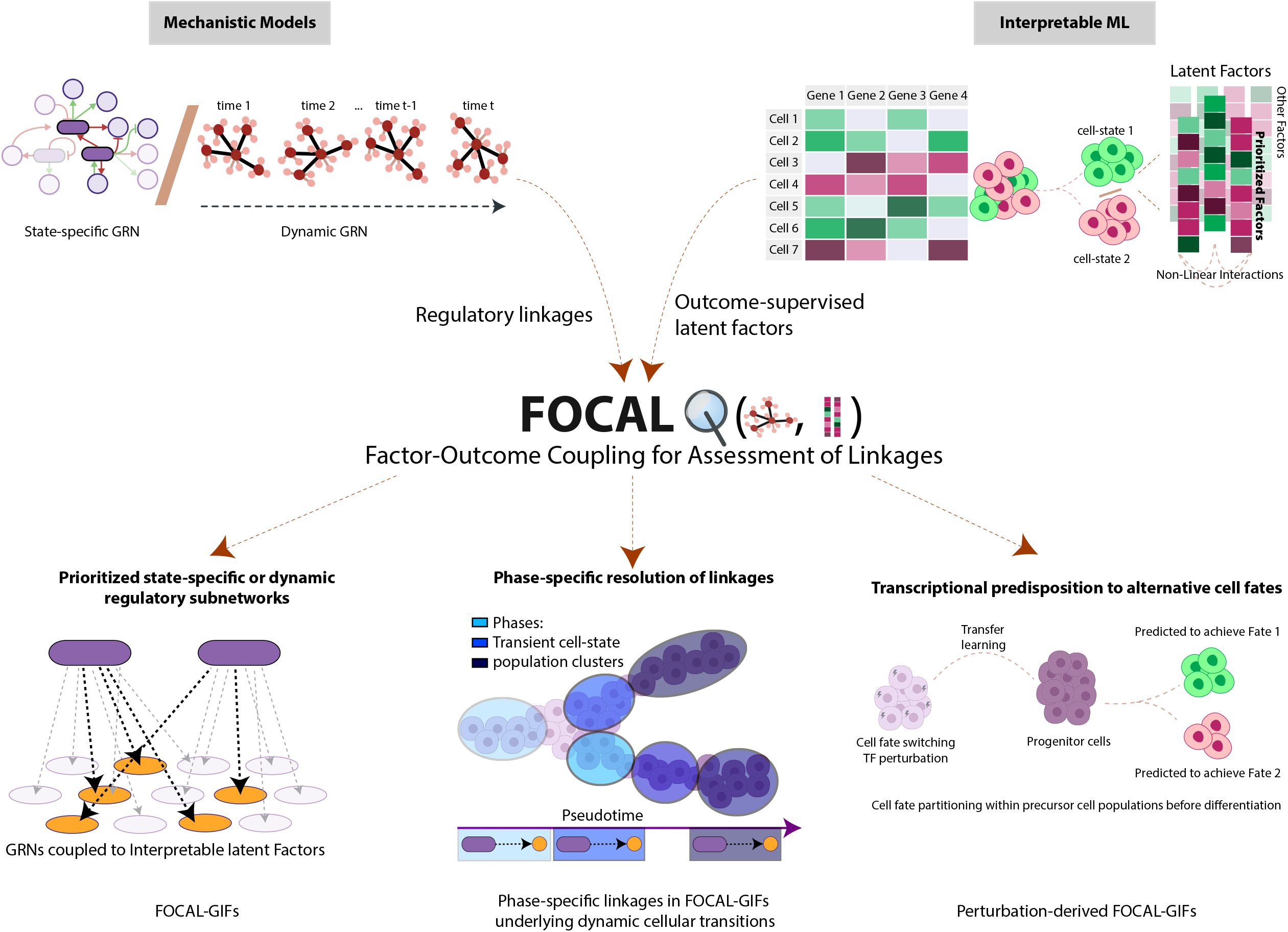
Overview of our FOCAL framework. Conceptual overview of FOCAL. The framework couples interpretable machine learning with mechanistic models (state-specific and dynamic GRNs) to dissect cell fate decisions. Regulatory linkages are supplied by the GRNs and outcome-associated latent factors by SLIDE, which is trained on the same transcriptomic data but without access to the network or its sequence priors. Their coupling yields three classes of outputs shown at the bottom: prioritized regulatory subnetworks (FOCAL-GIFs), temporal assignment of prioritized linkages to pseudo temporal phases, and prediction of transcriptional predisposition to alternative fates in uncommitted cells.

By projecting these outcome-associated latent factors onto the inferred GRNs, the coupled framework, which we term FOCAL (Factor–Outcome Coupling for Assessment of Linkages) prioritizes TF target-gene programs as well as regulatory linkages that are supported by both mechanistically informed network priors and information associated with the cellular-state contrasts. FOCAL denotes the framework; and in the present application to GRNs, the prioritized regulatory subnetworks generated by coupling LFs to GRNs are termed FOCAL-GIFs (GRNs coupled to Interpretable latent Factors). We emphasize that the candidate target gene sets that are tested are defined by the GRNs. Importantly, what is generated independently of these networks is the evidence used to evaluate them (**Fig. 1**). Applying this framework to B and T cell single cell trajectories, we successfully map stable-state transcriptional features to transient regulatory episodes that precede them and predict alternate fate predispositions in undifferentiated cells using TF perturbational datasets. The resulting FOCAL-GIFs uncovered an unanticipated cooperative interplay between the TFs NFATC2 and IRF8 in controlling B cell fate dynamics. Genetic perturbations were used to functionally validate this prediction, demonstrating that NFATC2 and IRF8 act cooperatively to restrain extrafollicular plasmablast differentiation and promote germinal center B cell fate during antigen-specific immune responses.

### Coupling interpretable ML to state-specific GRNs to prioritize regulatory subnetworks

To determine whether outcome-associated LFs could provide a distinct evidence layer for prioritizing regulatory programs within state-specific GRNs, we applied FOCAL to human B cell differentiation states. Upon activation, B cells can differentiate into extrafollicular plasmablast (PB) or germinal center (GC) states, which are regulated by counter-acting transcription factors. We previously assembled state-specific GRNs corresponding to the precursor GC (preGC) and PB states using an in vitro human B cell differentiation system and temporally resolved single-cell multiomics^15^. Here we asked whether projecting SLIDE-derived LFs, identified independently of network topology and sequence priors, onto these GRNs could delineate a prioritized set of regulatory subnetworks that discriminate the alternative states (**Fig. 2A**).

**Figure 2:**
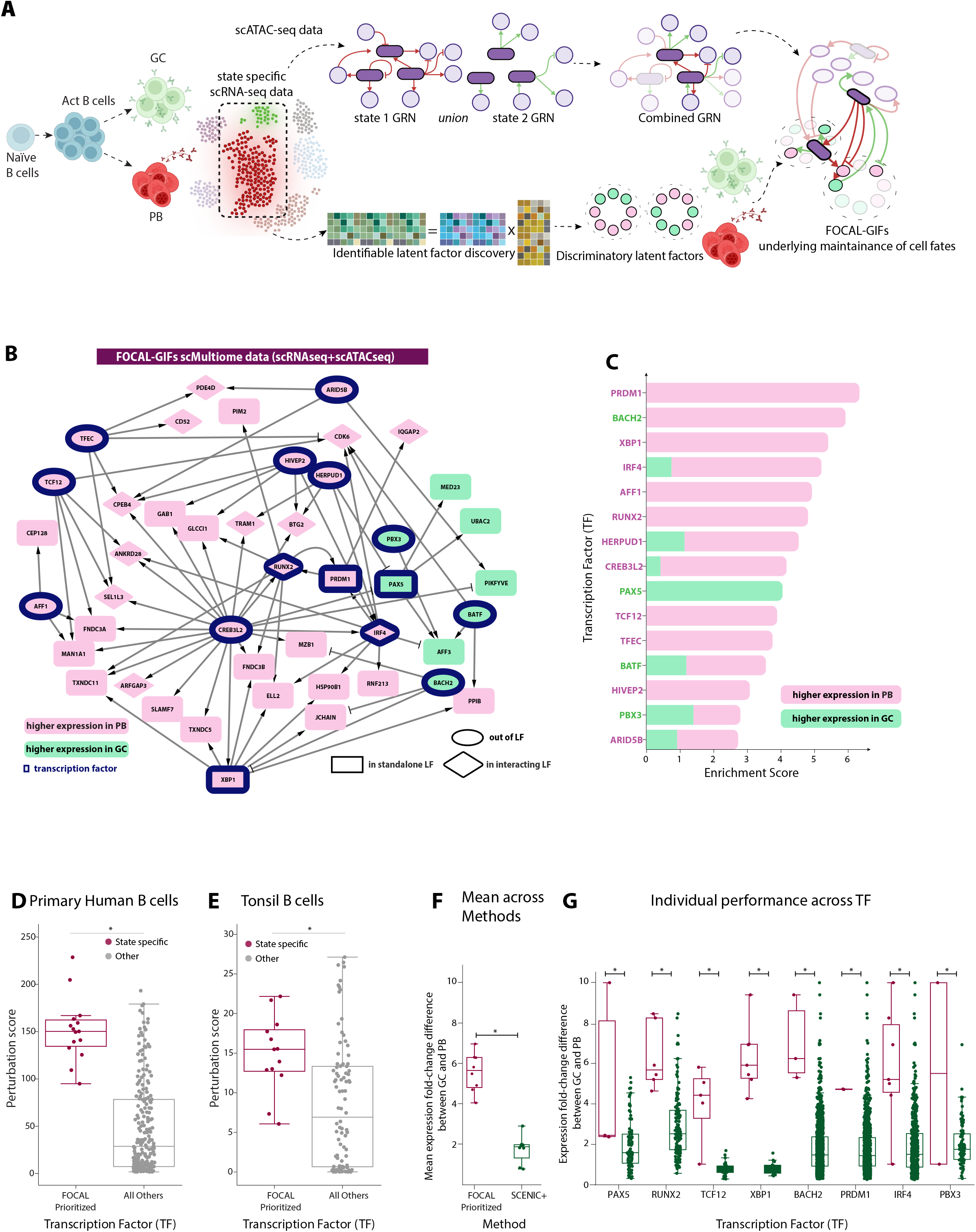
Coupling interpretable ML to state-specific GRNs to prioritize candidate regulatory subnetworks – FOCAL-GIFs. **A**. Conceptual overview of human B cell fate dynamics and coupling of complementary frameworks for the inference of state-specific GRNs and the identification of LFs that discriminate cell states using interpretable ML. **B**. Coupling SLIDE-derived LFs to state-specific GRNs identifies prioritized regulatory subnetworks (FOCAL-GIF). Colors indicate the state in which each gene is more highly expressed (green: higher expression in GC, pink: higher expression in PB). TFs are denoted by a blue boundary. TFs/genes in standalone and interacting LFs are denoted by rectangles and rhombuses respectively. TFs outside LFs are represented using oval shapes. **C**. Enrichment scores for the prioritized TFs from the state-specific GRNs. Colors indicate fraction of downstream genes for that TF with higher expression in the GC (green) or PB states (pink). **D**. Perturbation scores for FOCAL-prioritized TFs compared with all remaining TFs in the network, using *in silico* perturbations on state-specific GRNs in primary human B cells. **E**. Perturbation scores for the same prioritized TFs compared with all remaining TFs, using *in silico* perturbations on state-specific GRNs in tonsillar B cells. **F**. Distribution of mean absolute logFC of gene expression values of downstream target genes for indicated TFs shared between FOCAL and SCENIC+. Each point represents the average logFC (GC:PB) value for a TF across all the prioritized TF-gene links. **G**. The full distribution of absolute logFC values for downstream genes of each shared TF (PAX5, RUNX2, TCF12, XBP1, BACH2, PRDM1, IRF4, PBX3) as inferred by FOCAL and SCENIC+. Each point represents a TF-gene link-specific logFC (GC:PB) value.

Using SLIDE^14^ on the transcriptomic data derived from the multi-ome profiling, we first built a model encompassing 2 LFs (context-specific multi-gene co-expression programs) that could strongly discriminate the preGC and PB states (**Extended Figs. 1A, 1B**). We note that while the assembled GRNs corresponded to the preGC and PB states, they were reflective of GC and PB trajectories, and hence the terms preGC and GC are used accordingly in the manuscript. The SLIDE model was highly robust as evaluated using both k-fold cross-validation and phenotype-label permutation testing (**Extended Figs. 1A, 1B**). The identified state distinction generalized across multiome-derived RNA as well as scRNA-seq datasets (**Extended Figs. 1C-1E**). The two LFs were annotated using an LLM-based approach (**Extended Figs. 1F, 1G**).

We then tested every TF represented in the state-specific GRNs for enrichment of its downstream target genes within the two discriminatory LFs. The enrichment statistic accounted for both the size of each LF and the connectivity of the TF within the GRN (Methods). The two analyses contributed different forms of evidence to this integration: the GRNs were constructed from transcriptomic and chromatin-accessibility measurements together with sequence-based and TF-binding priors, whereas SLIDE identified LFs associated with the preGC-PB contrast without access to the inferred GRN or its sequence priors. Importantly, rather than simply ranking TF nodes, the integration prioritized a regulatory subnetwork comprising 15 TFs and 66 TF-gene linkages whose target genes were enriched within the discriminatory LFs (**Figs. 2B, 2C**).

Two consequences of the coupling are worth noting. First, prioritization did not require the gene encoding a TF to be a member of an LF; 10 of the 15 TFs anchoring this subnetwork were nominated entirely by the outcome-associated behavior of their downstream gene linkages, despite the transcript abundance of the TF itself being uninformative. The framework thereby provided a target-based readout of candidate TF activity. Second, targets assigned to individual TFs occurred within both the standalone and the interacting LFs (**Figs. 2B, 2C**). Because the LFs are themselves constructed from co-expression, this distribution is unlikely to reflect simple co-expression or redundant representations of a single transcriptional program. Instead, it is consistent with the SLIDE interaction term capturing higher-order relationships between partially distinct regulatory programs.

Having identified these specific regulatory linkages, we next asked whether they were constituted by TFs of higher regulatory importance within the large-scale human B cell GRN spanning various cell states. For this purpose, we used our previously developed perturbation score (PS)^15^, which estimates the predicted effect of propagating an insilico TF perturbation (computed using CellOracle^3^) through the GRNs underlying various cell states. This approach tests whether the outcome-associated prioritization identifies TFs that are also predicted to exert substantial influence within the GRN models. The LF-prioritized TFs had significantly higher PS values than the remaining TFs in the GRN (**Fig. 2D**). Thus, TFs selected using an evidence layer constructed without reference to network topology mapped preferentially onto nodes that the in-silico perturbation framework leveraging the GRN independently predicted to have larger perturbational consequences. The correspondence is notable in that it represents a convergence on TF nodes generated by two orthogonal modeling approaches that are based on complementary evidence layers.

We next asked whether this correspondence was retained when the same prioritized TFs were evaluated in GRNs assembled using an *ex vivo* tonsillar dataset^16^. Again, the same TFs (prioritized from the earlier analysis) showed significantly higher PS values than the remaining TFs (**Fig. 2E**). The reproduction of prioritized TFs using SLIDE and state-specific GRNs (constructed using *in-vitro* data) in independently derived tonsillar GC and PB networks indicates that the relationship between outcome-associated prioritization and predicted network influence is not restricted to the *in vitro* human B cell state GRNs within which the original TF nodes were prioritized.

We next evaluated the prioritized subnetworks, comparing them with regulons derived from SCENIC+^6^, a widely used method for GRN inference and corresponding subnetwork prioritization. The FOCAL-GIFs comprised far fewer TF-gene links, but the targets of these TF links showed substantially larger expression differences between the preGC and PB states than the targets within the much larger regulons prioritized by SCENIC+^6^ (**Figs. 2F, 2G**). Thus, the TF-gene linkages identified within FOCAL-GIFs are more specific and sensitive discriminators (mean expression fold difference) than the ones from SCENIC+ due to a fundamental conceptual difference. While SCENIC+ also prioritizes eRegulons using cell state labels analogous to FOCAL, it does so based on conventional univariate tests. These do not have the unique statistical guarantees – identifiability, inference and overall FDR control that SLIDE has^14^. Further, SLIDE does not leverage conventional network topological metrics. These have proved useful in prioritizing key TFs, but not in prioritizing key linkages as most prioritized TFs based on network topological metrics are high degree and centrality nodes and have a large set of linkages which cannot be further refined using topological metrics alone. SLIDE’s unique statistical properties have in the past enabled discovery of several novel cellular signaling and humoral mechanisms across contexts^14,17-20^. Here, we leverage the same statistical properties in a regulatory context to simultaneously prioritize both key TFs and linkages. In addition to novel TFs, the FOCAL-GIFs comprised major known regulators of PB differentiation, including IRF4, PRDM1 and XBP1, as well as those regulating GC B cell trajectory, namely PAX5, BACH2 and BATF (**Figs. 2B, C**). The recovery of subnetworks involving major lineage-defining TFs for the PB and GC trajectories provided strong biological support for the prioritization strategy for the FOCAL framework..

### Prioritization of regulatory subnetworks generalizes to closely related cell states

The B cell analyses above are reflective of two states that differ substantially in their transcriptional programs. To test whether the framework generalizes to a more nuanced discrimination task and to a different cellular context, we applied it to terminally exhausted CD8 T cells (Tex^term^) and their phenotypically variant cytotoxic counterparts expressing the killer cell lectin-like receptor (Tex^KLR^) (**Extended Fig. 2A**)^21^.

A SLIDE model provided significant discrimination between Tex^term^ and Tex^KLR^ CD8 T cells (**Extended Fig. 2B**) and comprised two standalone and two interacting LFs (**Extended Fig. 2C**). We constructed state-specific GRNs for the two states and coupled these to the SLIDE-derived LFs as described above. Coupling resolved the large scale GRNs into a prioritized regulatory subnetwork comprising 40 candidate TF-target linkages anchored by 14 TFs (**Extended Fig. 2D**). As in the B cell analysis, the target genes included those preferentially expressed in either state (**Extended Fig. 2E**), and genes downstream of the same TF spanned both the standalone and the interacting LFs (**Extended Fig. 2D**), consistent with the SLIDE interaction term capturing higherorder regulatory relationships.

The TFs driving these prioritized linkages exhibited significantly higher perturbation scores than the remaining TFs in the comprehensive GRNs (**Extended Fig. 2F**), reproducing the correspondence between outcome-associated prioritization and model-predicted network influence in closely related alternative cellular states. Comparison with SCENIC+^6^ again showed that the prioritized subnetworks comprised a small number of TF-gene linkages with significantly better sensitivity and specificity than the regulons recovered by SCENIC+ (**Extended Figs. 2G, 2H**). The prioritized subnetwork comprised novel candidates as well as key known TFs and genes underlying CD8 exhaustion including Batf, Nfatc1, Foxo1, Tbx21, Bhlhe40, Pdcd1 and Tox (**Extended Fig. 2D**)^22,23^. Thus, the analysis provided strong biological support for the prioritization strategy when applied to closely related cell states. Altogether these analyses indicate that coupling outcome-associated latent factors with inferred GRNs can prioritize regulatory subnetworks, that are biologically supported, across highly distinct as well as closely related cell states.

### Coupling interpretable ML to dynamic GRNs resolves specific regulatory linkages driving cell state transitions

While state-specific GRNs are powerful and widely used representations of gene regulatory architectures, they preferentially capture TF-target gene relationships that are reflective of stable or prevalent cellular states. Such GRNs have limitations when used to analyze cell-fate transitions. TFs that function dynamically at a cell fate bifurcation or within transition states in a differentiation trajectory can exert substantial regulatory impact during such transitions without being required for the maintenance of stable regulatory states. Such dynamic TF activity may therefore be poorly represented in GRNs assembled from cells occupying discrete or relatively stable states. Dynamic GRN models address this problem by reconstructing changes in regulatory relationships along differentiation trajectories rather than representing them only within defined cellular states (**Fig. 3A**)^4,7^.

**Figure 3:**
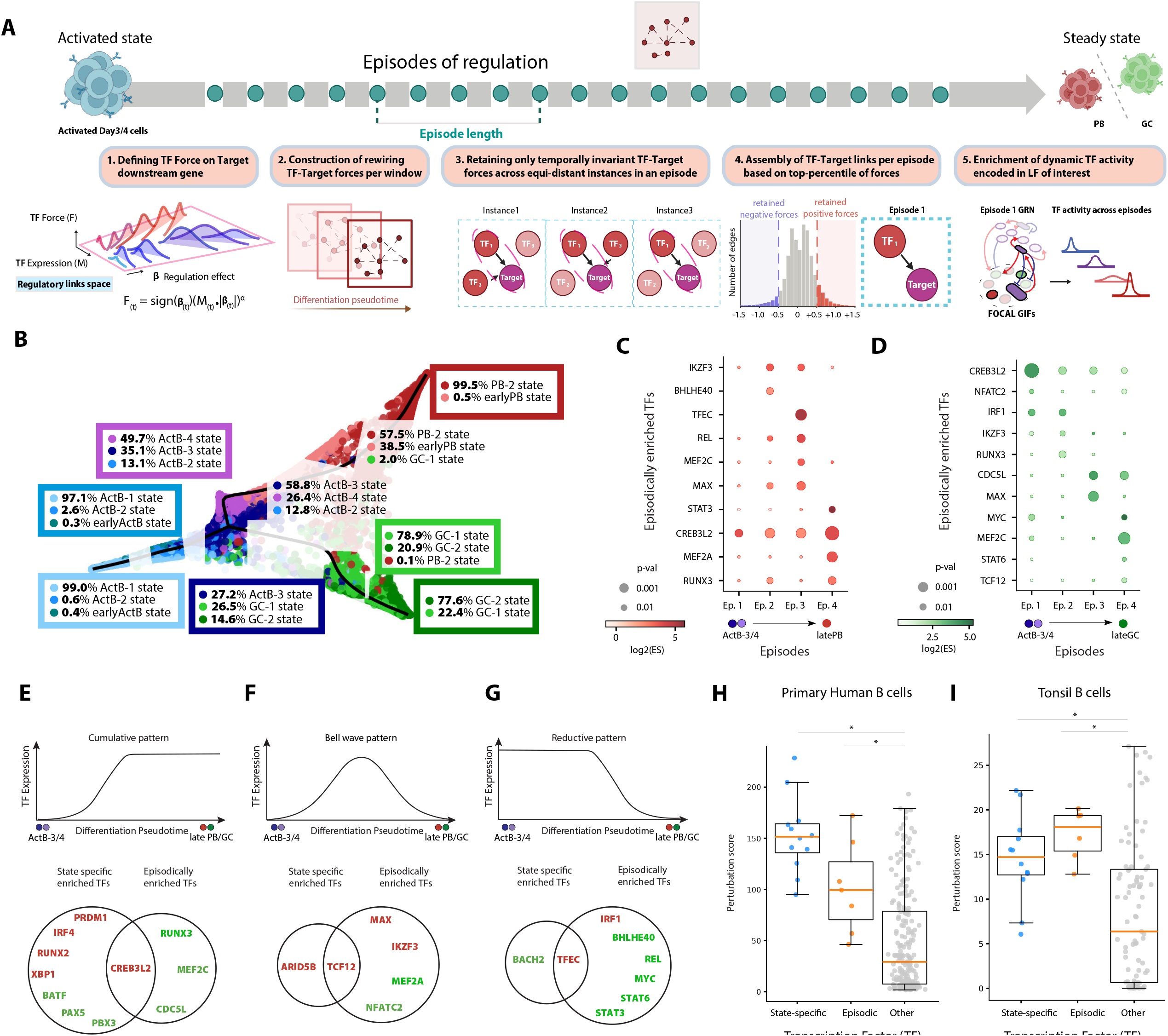
Coupling interpretable ML to dynamic GRNs resolves specific regulatory linkages driving cell state transitions. **A**. Defining episodes of TF activity. The force landscape of a TF regulating a target gene is defined across the trajectory in episodes. Invariant TF-Target forces are retained to construct episodic GRNs, temporally quantifying the regulatory action of TFs. **B**. Population changes across the B-cell differentiation trajectory over 7 days, along with the split between cell-state proportions within a window, are displayed for exemplary sampled windows. Black line indicates the fitted curve, with pre- and post-bifurcation branching points marked on the trajectory. **C, D**. Enrichment scores for the prioritized TFs from the dynamic GRNs that exert influence episodically across the PB (C) and GC (D) branches. Dot color intensity indicates enrichment effect size, and dot size indicates statistical significance. **E, F, G**. Overlap between state-specific and episodically enriched TFs based on classification of their expression patterns along the trajectory: Cumulative, monotonically increasing (E); Bell-shaped, transiently increasing (F); and Reductive, monotonically decreasing (G). Venn diagrams show TFs prioritized from state-specific GRNs, from dynamic GRNs, or from both. **H, I**. Comparison of perturbation scores for prioritized TFs from the state-specific and dynamic analyses vs all remaining TFs. Perturbation scores computed using GRNs derived from primary human B cells (H) or tonsillar B cells (I), as in Fig. 2D.

We reasoned that transiently acting TFs could nevertheless contribute to establishing downstream gene programs that persist in the absence of the TF and thereby could be captured within an outcome-associated LF. In this sense, the transcriptional programs of established cellular states can retain information about regulatory events that occurred during their formation. Projection of these LF captured programs onto dynamic GRNs therefore provides a means of retrospectively associating stable fate-associated transcriptional features with candidate regulators acting during preceding transition states or fate bifurcations (**Fig. 3A**).

We therefore tested whether the same outcome-associated SLIDE LFs used to prioritize the state-specific GRNs could be projected onto temporally resolved GRNs to reveal regulatory subnetworks operating during intervals of human B cell differentiation. Using Dictys^4^, we reconstructed dynamic GRNs along trajectories from activated B cells toward the PB and preGC states (**Fig. 3B**) based on a previously established pseudotime trajectory^15^. To couple the Dictys-derived networks with the SLIDE-derived LFs, we extended the dynamical analysis to define regulatory episodes along differentiation pseudotime. Three temporal units are used throughout. A window is the smallest time interval over which a TF-target relationship is estimated; an episode is a run of equidistant windows on the trajectory, across which that relationship remains temporally invariant; and a phase, introduced below, is a contiguous block of time over which the peaks of dynamic TF activity of prioritized links are empirically observed. Episodes and phases are therefore both aggregates of cell-state over time, distinguished by whether they are defined by regulatory invariance or by cell-state composition. We estimated TF effects on downstream genes across successive windows using the TF-force metric, which combines TF expression with the inferred sign and strength of its regulatory effect on a target gene. Regulatory linkages with consistent positive or negative effects within an episode were retained and the strongest of these linkages were assembled into episode-specific GRNs. (**Figs. 3A and 3B**, Methods). These episodic networks were then tested for enrichment of TF target genes within the same SLIDE-derived LFs used in the state-specific analysis. Thus, the outcome-associated evidence layer was held constant while the regulatory representation was changed from state-specific to dynamic GRNs.

This analysis resolved a dynamic regulatory subnetwork comprising 62 specific temporal linkages, anchored by 16 TFs, whose downstream genes were enriched within the LFs discriminating the preGC and PB states (**Figs. 3C and 3D, Extended Figs. 3A-3G)**. Importantly, these regulatory linkages were identified within specific regulatory episodes and did not necessarily persist into the terminal PB or preGC states. The linkages defining this subnetwork were driven by TFs associated with both PB-and GC-differentiation, that were predicted to function at distinct positions along the two trajectories (**Figs. 3C and 3D, Extended Figs. 3A-3G**). Comparison with TF expression dynamics revealed three broad temporal patterns: TFs whose expression increased cumulatively toward the terminal states (**Fig. 3E**), regulators that were transiently expressed during differentiation (**Fig. 3F**), and regulators whose expression diminished as differentiation progressed (**Fig. 3G**). The prioritized TFs and corresponding linkages in the dynamic GRN corresponded to all three kinds of patterns and were mostly distinct from the TFs prioritized from the state-specific GRNs (**Figs 3E-3G**). These analyses demonstrated that the GIFs obtained by coupling SLIDE LFs with dynamic GRNs can reveal candidate regulatory programs associated with temporally resolved intervals within differentiation trajectories, encompassing transition states or fate bifurcations, that are not readily resolved using state-specific GRNs.

We next asked whether the TFs anchoring these transient, phase-specific linkages were also predicted to exert substantial global influence within the larger GRNs. Importantly, TFs prioritized from the FOCAL-GIFs obtained by coupling dynamic GRNs to SLIDE exhibited PS values that were significantly higher than perturbation scores for TFs not prioritized by the FOCAL-GIFs (**Figs. 3H and 3I**). The former PS values were comparable to those for TFs prioritized by FOCAL-GIFs via coupling of state-specific GRNs to SLIDE. These results remained consistent irrespective of whether the PS scores were computed using propagation along GRNs inferred from our primary B cell system (**Fig. 3H**) or tonsil-derived B cell data (**Fig. 3I**). We note that the TFs from the FOCAL-GIFs also had higher PS values than those prioritized by Dictys (**Extended Figs. 3H and 3I**). These results indicate that the same outcome-associated LFs can be projected onto both state-specific and dynamic regulatory GRNs to prioritize largely non-overlapping sets of TFs, only CREB3L2 and TFEC being recovered by both, that appear to act complementarily in regulating cell fate transitions and the maintenance of stable differentiated states.

### Latent-factor-prioritized TF–target relationships resolve phase-specific regulatory activity

The preceding analyses showed that outcome-associated LF enrichment prioritizes TFs that both state-specific and dynamic GRNs predict to have substantial regulatory influence. We next asked whether prioritization could be resolved beyond individual TFs to specific TF-target relationships, and whether these could be located along the differentiation trajectories. For each candidate TF-target linkage we determined the maximal TF-force, as defined above, across a given trajectory. LF-prioritized TF-target relationships retained within both the state-specific and dynamic GRN subnetworks exhibited significantly greater regulatory-force magnitudes than the remaining GRN edges (**Fig. 4A**). Thus, the prioritized linkages occupy the high-force regions of the inferred regulatory landscape rather than being distributed randomly across it.

**Figure 4:**
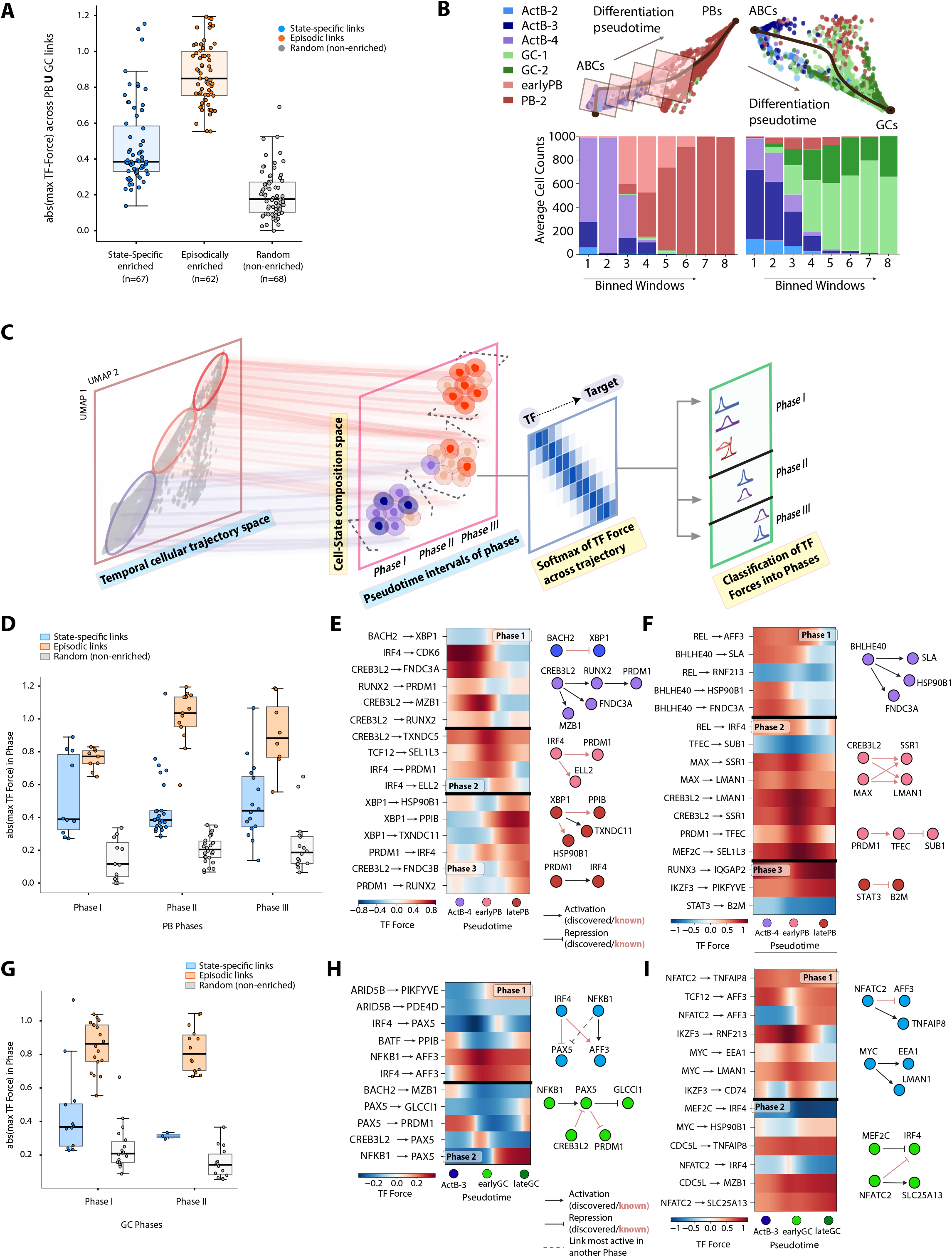
FOCAL-GIF TF-target linkages resolve phase-specific regulatory activity. **A**. Evaluation of regulatory activity (quantified using absolute value of the maximal TF force) for the prioritized TF-gene links (from the state-specific and dynamic GRN analyses) as well as randomly sampled non-enriched links across all phases of regulatory activity in both branches. **B**. Capturing cellular heterogeneity during differentiation by binning windows across the trajectory. Sliding windows are binned into eight bins per branch to display cell-state proportions. **C**. Temporal clustering of TF forces regulating state transitions in phases. Cell-state transitions on each branch occur in phases, which represent shifts in majority state population on the trajectory. Phase switches are quantified by the majority state’s termination pseudotime, indicating loss of that state’s identity coincident with a switch in transcriptional regulatory activity. **D**. Evaluation of regulatory activity (quantified using TF force) for the prioritized TF-gene links (from the state-specific and dynamic GRN analyses) as well as randomly sampled non-enriched links in a phase-specific fashion for the PB branch. **E**. Visualization of regulatory activity of key TF-gene links identified from the statespecific GRN analysis across three phases in the PB branch. Heatmap shows TF force along pseudotime; network diagrams at right summarize activating and repressive linkages, distinguishing those newly discovered here from those previously known, as keyed. **F**. Visualization of regulatory activity of key TF-gene links identified from the dynamic GRN analysis across three phases in the PB branch. Display as in E. **G**. valuation of regulatory activity (quantified using TF force) for the prioritized TF-gene links (from the state-specific and dynamic GRN analyses) as well as randomly sampled non-enriched links in a phase-specific fashion for the GC branch. **H**. Visualization of regulatory activity of key TF-gene links identified from the statespecific GRN analysis across both phases in the GC branch. Display as in E; dashed edges denote links whose maximal force falls in another phase. **I**. Visualization of regulatory activity of key TF-gene links identified from the dynamic GRN analysis across both phases in the GC branch. Display as in E.

We next asked how the prioritized relationships were distributed temporally during the cell-fate transitions. Dynamic TF activity was resolved into successive windows that were grouped into discrete phases along the PB and GC trajectories (**Fig. 4B**), and each prioritized TF-target relationship was assigned to the phase in which its regulatory-force magnitude was maximal (**Fig. 4C**). This converted the prioritized subnetwork into a temporally ordered representation of regulatory activity. Three phases of regulatory activity were resolved along the PB trajectory (**Extended Fig. 4A)**. Within each phase, TF-target relationships prioritized from both the state-specific and the dynamic GRNs exhibited greater regulatory-force magnitudes than comparator edges in the corresponding network (**Fig. 4D**). Independent support was provided by TF-binding inference, which is external to the regulatory-force metric. Consistent with expectation, inferred binding for TFs prioritized from both the state-specific and dynamic GRNs was higher than for the remaining TFs across all three phases (**Extended Fig. 4B**). Key regulatory relationships from the state-specific (**Fig. 4E**) and the dynamic GRNs (**Fig. 4F**) corresponded to both known for example, IRF4→PRDM1, ELL2 in phase 2 and PRDM1→IRF4 in phase 3 and novel regulatory relationships. The expression and inferred regulatory dynamics of selected TFs, namely BACH2, IRF4 and PRDM1 were consistent with their established behavior during PB differentiation (**Extended Figs. 4C, 4D**).

A corresponding analysis of the GC trajectory resolved two phases of regulatory activity (**Extended Fig. 4E**). As observed for the PB branch, TF-target relationships prioritized from either state-specific or dynamic GRNs showed greater regulatory-force magnitudes than the remaining edges within both phases (**Fig. 4G**). The prioritized links from both the state-specific (**Fig. 4H**) and dynamic GRNs (**Fig. 4I**) captured known and novel activating and repressive relationships. As earlier, inferred binding for TFs prioritized from both the state-specific and dynamic GRNs was higher than for the remaining TFs across both phases (**Extended Fig. 4F**). The expression and regulatory dynamics of representative TFs were consistent with established patterns during GC differentiation (**Extended Figs. 4G, 4H**).

Together, these analyses extend LF-based prioritization from regulatory nodes to individual candidate TF-target linkages and locate these within successive waves of regulatory activity during differentiation. Conventional network-topological measures such as degree or centrality rank TFs according to their connectivity within the complete GRN, but do not specify which of a TF’s many target linkages are most closely associated with the phenotypic contrast, or when along a differentiation trajectory those linkages become prominent. By integrating outcome-associated LFs with state-specific and dynamic GRNs, the framework resolves a restricted set of candidate regulatory edges and organizes them according to their temporal dynamics during cell-fate transitions.

### Perturbation-derived latent factors reveal transcriptional predisposition to alternative cell fates

Having shown that outcome-associated LFs can prioritize regulatory programs within stable cellular states and along differentiation trajectories, we next asked whether LFs learned from fate-altering TF perturbations could identify transcriptional predisposition to alternative cell fates in progenitors before overt lineage divergence. This represents a conceptual next step in the framework. Rather than learning cellular programs from the differentiated GC and PB states, we sought to learn programs induced by perturbations of fate-determining TFs and then project these programs onto unperturbed activated B cells that represent progenitors of GC and PB cells (**Fig. 5A**).

**Figure 5:**
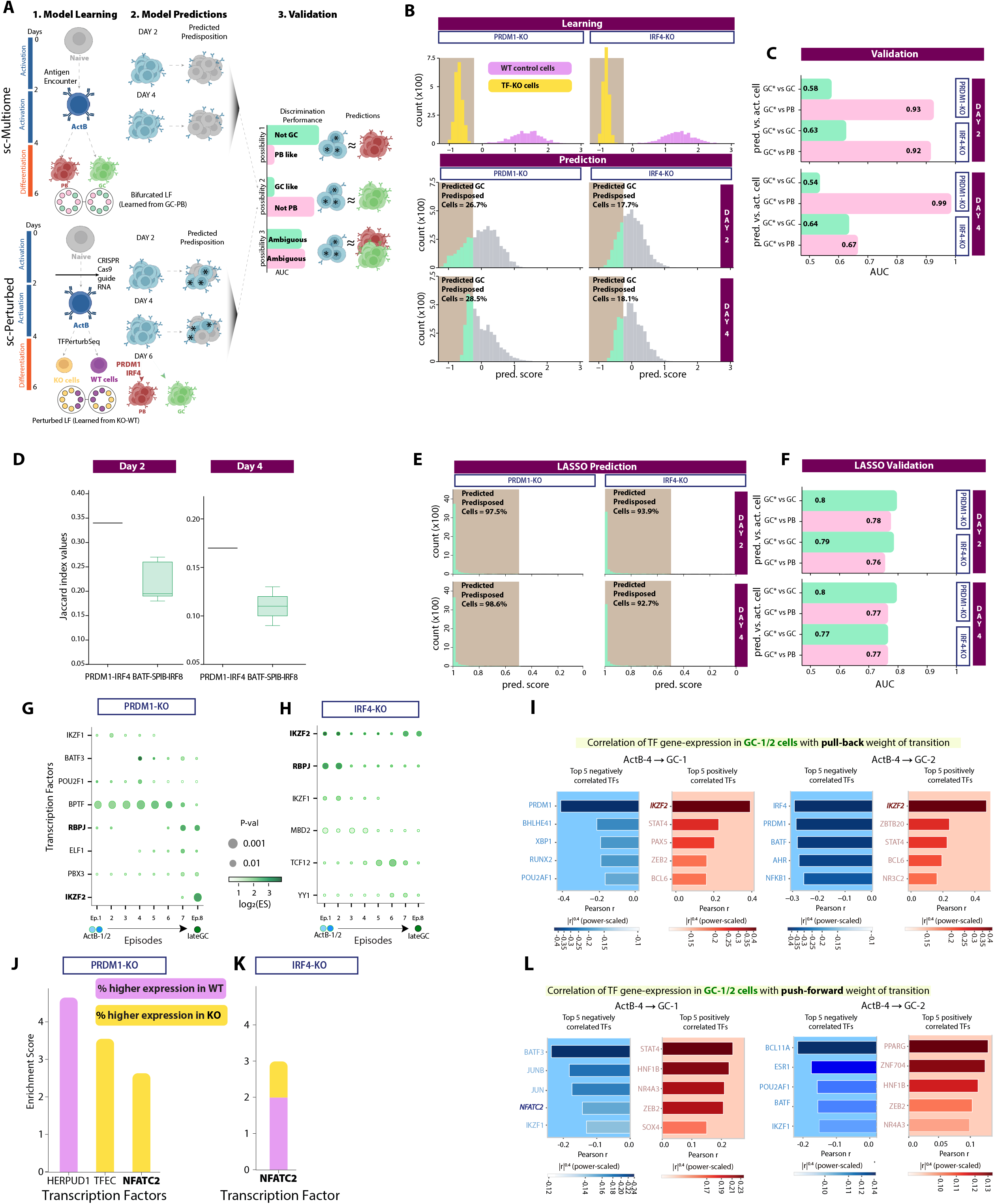
Perturbation-derived LFs reveal transcriptional predisposition to alternative cell fates. **A**. LFs learned from bifurcated states in the upper panel and from TF knockout versus unperturbed control cells in the bottom panel are projected onto early ABCs at Day 2 and Day 4 using the multi-ome dataset to determine whether these programs can predict cell fate in progenitor compartment. Validation is performed by quantifying the fraction of progenitor cells that are assigned as predisposed to one or the other fate (PB or preGC) with an assessment of their transcriptional similarity to corresponding bifurcated states. **B**. Projection of LFs learned from the PRDM1-KO and IRF4-KO experiments onto progenitor populations i.e., unperturbed ABCs at day 2 and day 4. Top panels visualize distribution of LF scores for PRDM1-KO cells (yellow) vs unperturbed (purple) cells and IRF4-KO (yellow) versus unperturbed cells (purple). Projecting the scores on ABCs at Day 2 and Day 4 (bottom panels) highlights a subset of ABCs (green portion) exhibiting early GC-like fate predisposition at the transcriptional level. **C**. Performance of the SLIDE model in discriminating predicted GC predisposed cells within the progenitor population (ABCs at Days 2 and 4) against bifurcated GC or PB cell states. **D**. Overlap between the sets of progenitor cells independently predicted as GC-predisposed by the IRF4-KO and PRDM1-KO models, compared with the overlap expected from predictions based on unrelated TF perturbations. **E**. Projection of a LASSO model learned from the PRDM1-KO and IRF4-KO experiments onto progenitor populations i.e., unperturbed ABCs at day 2 and day 4. Projecting the scores on ABCs at Day 2 and Day 4 (bottom panels) highlights a subset of ABCs (green portion) exhibiting early GC-like fate predisposition at the transcriptional level. **F**. Performance of the LASSO model in discriminating predicted GC predisposed cells within the progenitor population (ABCs at Days 2 and 4) against bifurcated GC or PB cell states. **G, H**. TFs whose downstream genes are episodically enriched in the LFs discriminating PRDM1-KO (G) and IRF4-KO (H) versus unperturbed cells. Color scale indicates enrichment effect size (log_2_ ES) and dot size indicates statistical significance. **I**. TFs positively or negatively regulating transitions from Day 4 ABCs to GC B cells learned using an optimal transport-based model leveraging the pull-back distribution. **J, K**. Enrichment scores for the prioritized TFs from the PRDM1-KO (J) and IRF4-KO (K) experiments. Colors indicate higher expression in control (purple) or KO (yellow) states. **L**. TFs positively or negatively regulating transitions from Day 4 ABCs to GC B cells learned using an optimal transport-based model leveraging the push-forward distribution.

We first asked whether the SLIDE-derived LFs that discriminate established GC and PB states were sufficient to identify transcriptional divergence in activated B cells suggestive of fate divergence. Although the standalone LF strongly discriminated the differentiated GC and PB states, neither the standalone or the inclusion of interacting LFs reliably resolved subsets of activated B cells (day 2 or day 4) and predicted their predisposition toward either fate (**Extended Figs. 5A and 5B**). Thus, transcriptional programs that discriminate established states do not necessarily contain sufficient information to identify early cells biased toward those states in the progenitor compartment.

We therefore reasoned that perturbations of lineage-determining TFs could provide a more direct means of learning transcriptional programs associated with fate predisposition (**Fig. 5A**). We focused on learning from Perturb-seq experiments corresponding to the knockouts of IRF4 and PRDM1, two critical regulators of PB differentiation whose loss results in blocks to PB differentiation and promotes generation of GC precursors^15^ (**Fig. 5A**). Using SLIDE on the transcriptomic data for both the IRF4 and PRDM1 KO experiments, we were able to identify LFs (one significant LF for each TF perturbation using corresponding SLIDE model) that provided excellent discrimination between the perturbed and unperturbed states (**Figs. 5B, Extended Fig. 5C and 5D**). These IRF4 or PRDM1 perturbation-associated LFs were then projected onto unperturbed day 2 and day 4 activated B cells to identify cells in the progenitor compartment whose transcriptional profiles were aligned with the TF-KO state and were therefore predicted to be GC-predisposed (**Fig. 5B**).

We next assessed whether these predicted transcriptionally predisposed cells manifested features of their downstream GC state. In this test, near-chance discrimination (AUC close to 0.5) indicates transcriptional similarity to the reference state and is therefore the expected outcome of a correct prediction, whereas strong discrimination indicates dissimilarity. When the predicted subsets of unperturbed early and late activated B cells were compared to bifurcated GC cells, they were almost indistinguishable (**Fig. 5C**). Further, they were strongly distinguishable from PB cells (**Fig. 5C**), consistent with acquisition of a GC-associated transcriptional bias preceding their overt differentiation. The strong discrimination from PB cells in the same comparison provides the corresponding positive control, establishing that the classifier had sufficient power to separate cells of the two fates when they differ. Although independent SLIDE models were built from the two perturbation experiments, the cells predicted to be predisposed by each model overlapped significantly, consistent with the convergent roles of these two TFs in promoting the alternative PB differentiation program (**Fig. 5D**). While these analyses do not establish the eventual fates of individual cells in the progenitor compartment, they resolve transcriptional states consistent with early, dynamic programming of alternative fates. Together, these results demonstrate that the SLIDE-derived LFs learn aspects of higher-order dynamic regulatory activity underlying cell fate decisions that can then be used to make credible fate predictions for unperturbed cells before they have undergone lineage specification.

We compared the SLIDE-based approach with two alternative strategies trained on the same perturbation datasets. LASSO regression assigned nearly all early activated cells to the predisposed class and failed to reproduce the close transcriptional correspondence between the predicted cells and the GC reference state (**Figs. 5E and 5F**). An alternative model based on conventionally defined differentially expressed genes similarly failed to resolve comparably informative subsets (**Extended Figs. 5E and 5F**). Thus, the perturbation-derived LFs, which inherit the statistical properties of the SLIDE model, contain information about transcriptionally predisposed states that is not equivalently recovered by gene-level or alternative multivariate approaches.

As before, we sought to test whether the FOCAL framework generalizes to learning from more nuanced TF perturbations in a different cellular context. We applied it to precursor exhausted CD8 T (T_pex_) cells capable of responding to existing immunotherapies as well as terminally exhausted T (T_ex_) cells not capable of responding to cancer immunotherapy (**Extended Fig. 6A**)^24^. Prior work has implicated a key role for IKZF1 in driving T_pex_ cells away from to the T_pex_ to the T_ex_ state^24^. Given this, we hypothesized that IKZF1 KO would restore cells towards a T_pex_ state and we would be able to learn the corresponding program using our framework. Using SLIDE on the transcriptomic data for the IKZF1 KO experiment, we were able to identify one significant LF that provided excellent discrimination between the perturbed and unperturbed states (**Extended Figs. 6B and 6C**). As earlier, the perturbation-associated LF was then projected onto T_ex_ cells to identify a subset whose transcriptional profile was aligned with the TF-KO state and were therefore predicted to be more like the T_pex_ state (**Extended Fig. 6D**). Indeed, when the subset of T_ex_ cells predicted to be T_pex_ like were compared to true T_pex_ cells, they were more transcriptionally similar to these T_pex_ cells than cells not predicted to be predisposed (**Extended Fig. 6E**). The discrimination achieved by our framework significantly surpassed the discrimination achieved by LASSO or DE analyses (**Extended Figs. 6F-6H**). Together, these results demonstrate that using the SLIDE-derived LFs, one can learn how perturbations of lineage-determining TFs in multiple systems define transcriptional programs associated with fate predisposition.

**Figure 6:**
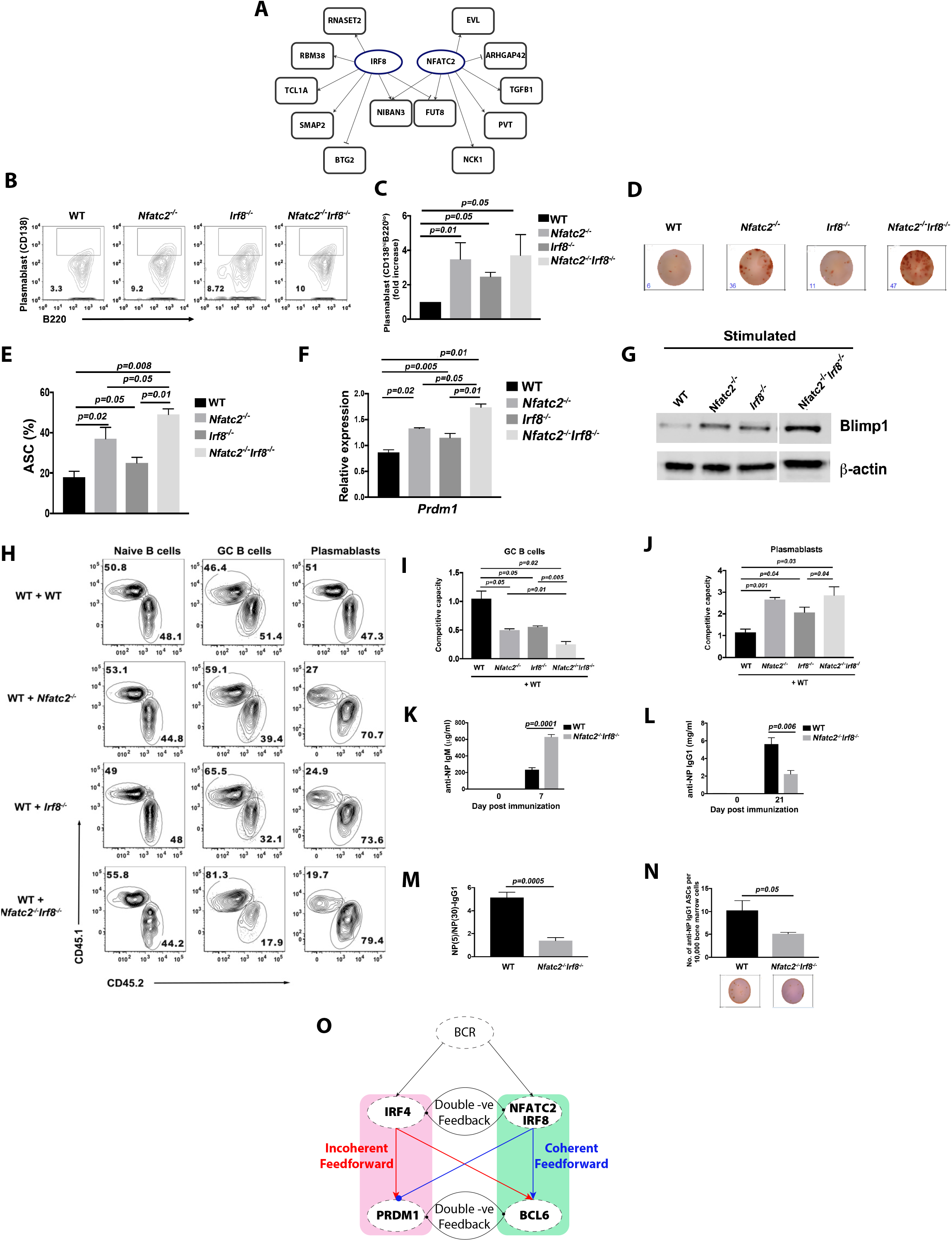
NFATC2 acts cooperatively with IRF8 to restrain plasmablast differentiation and promote the germinal center fate. **A**. Sub-GRN comprising the genes assigned downstream of NFATC2 and IRF8 in the inferred networks. NIBAN3 and FUT8 are targeted by both factors. The remaining genes are specific to either NFATC2 or IRF8. **B**. Representative flow cytometry plots demonstrating frequency of plasmablasts in WT, *Nfatc2*−/−, *Irf8*−/− and *Nfatc2*−/−*Irf8*−/− in anti-IgM, anti-CD40, IL-2,-4,-5 stimulated B cell cultures. In vitro differentiation of B cells was performed for 3 days. **C**. Quantification of frequency of plasmablasts corresponding to the flow cytometry plots in (B). Quantified as fold increase in CD138+B220lo plasmablasts relative to WT cultures. Bars show mean and SEM. **D**. Representative ELISPOT wells for WT, *Nfatc2*−/−, *Irf8*−/− and *Nfatc2*−/−*Irf8*−/− B cells. Numbers below each well indicate spot counts. **E**. Frequency of antibody secreting cells within WT, *Nfatc2*−/−, *Irf8*−/− and *Nfatc2*−/−*Irf8*−/− B cells. Quantified from the ELISPOT assay in (D). Bars show mean and SEM. ***F***. *Prdm1* transcript levels in *Nfatc2*−/−, *Irf8*−/− and *Nfatc2*−/−*Irf8*−/− B cells, measured by RT-qPCR, normalized to [housekeeping gene] and expressed relative to WT. Bars show mean and SEM. **G**. Blimp1 protein levels (encoded by the *Prdm1* gene) in *Nfatc2*−/−, *Irf8*−/− and *Nfatc2*−/− *Irf8*−/− B cells. Beta-actin serves as the loading control. **H**. Representative flow cytometry plots from competitive adoptive transfers of naive B cells, GC B cells and plasmablasts recovered from recipient mice co-transferred with WT cells (CD45.1 mice) and WT, *Nfatc2*−/−, *Irf8*−/− or *Nfatc2*−/−*Irf8*−/− cells (CD45.2 mice). Numbers correspond to the frequency of each population, analyzed 14 days after immunization with NP-KLH. **I, J**. Quantification of GC B cells (I) and plasmablasts (J) corresponding to the competitive adoptive transfer experiments in (H), expressed as competitive capacity relative to the co-transferred WT reference population. Bars show mean and SEM. **K, L**. NP specific serum antibody responses after immunization. Anti-NP IgM at days 0 and 7 (K) and anti-NP IgG1 at days 0 and 21 (L) in WT and *Nfatc2*−/−*Irf8*−/− mice measured by ELISA. **M**. Ratio of high affinity (NP5) to total (NP30) anti-NP IgG1 in WT and *Nfatc2*−/−*Irf8*−/− mice, quantifying affinity maturation, measured at day 21. **N**. Number of anti-NP IgG1 ASCs per 10,000 bone marrow cells, with representative ELISPOT wells shown below each bar. **O**. Updated mechanistic model demonstrating co-operative action of NFATC2 with IRF8 in promoting the GC fate and restraining plasmablast differentiation. NFATC2 and IRF8 constitute a coherent feed-forward loop downstream of BCR signaling that opposes the incoherent IRF4-PRDM1 loop.

### Contextualizing perturbation-derived latent factors using state-specific and dynamic GRNs

We next sought to interrogate the learnt LFs that reflected GC predisposition with previously assembled B cell state-specific and dynamic GRNs. This reflects a conceptual parallel to the prior analyses where the coupled interrogation of SLIDE-derived LFs and GRNs provides novel insights. Projection of the perturbation-derived LFs onto the dynamic GRNs identified phase-specific enrichment of target programs for several TFs along the GC differentiation trajectory (**Figs. 5G and 5H**). Interestingly, two TFs – IKZF2 (Helios) and RBPJ were shared across the two independent analyses. Although the role of IKZF2 is well characterized in terms of controlling developmental trajectories for T cells^25,26^, it has an underappreciated role in the formation of GC B cells^27^. The inferred regulatory activity of IKZF2 was higher than many known GC lineage-defining TFs which had higher expression. The functioning of IKZF2 in the GC differentiation trajectory was supported by an alternative multimodal optimal-transport analysis^28^ which also recovered well known lineage-defining TFs (**Fig. 5I**). Together, this provided additional support for the association of IKZF2 with the GC-directed regulatory program.

Projection of the same perturbation-derived LFs onto state-specific GRNs identified target programs for three TFs enriched in the PRDM1-KO LF and two TFs enriched in the IRF4-KO LF (**Figs. 5J and 5K**). Strikingly, NFATC2 was prioritized in both analyses and in episodic-enrichment of outcome-derived LFs in the GC branch (**Figs. 3D and 3F**). Like IKZF2, the inferred regulatory activity of NFATC2 was higher than many known GC lineage-defining TFs which had higher expression. This inference was also supported by the multimodal OT^28^ framework (**Fig. 5L**). Thus, two distinct perturbations that bias cells away from the PB trajectory converged on an inferred NFATC2 regulatory program, nominating NFATC2 as a new candidate regulator of the GC-PB fate decision.

### Functional validation of NFATC2 in regulating GC fate with IRF8

We next sought to functionally test the regulatory role of NFATC2 predicted by the coupled perturbation-LF and GRN analyses. The GC-PB fate bifurcation is controlled by sequentially acting, mutually antagonistic regulatory modules in which the TFs, IRF8 and BCL6 promote GC differentiation, whereas IRF4 and PRDM1 promote PB differentiation^29-31^. Because loss of either IRF4 or PRDM1 promotes differentiation toward a GC-associated state^29-31^, we asked whether the perturbation-derived LFs contained target programs shared with established GC-promoting regulators. The LFs derived from both perturbations were enriched for NFATC2 target genes and contained a subset of genes assigned as targets of both NFATC2 and IRF8 in the inferred GRNs (**Fig. 6A**). This convergence suggested that NFATC2 and IRF8 might act cooperatively within a GC-associated regulatory program and in so doing antagonize the PB fate.

Given the evolutionary conservation of the core B cell regulatory modules, we hypothesized that this newly predicted NFATC2/IRF8 cooperative interplay would be functionally conserved in mice. Utilization of a murine model system allowed us to rigorously test the inferred regulatory relationship and its impact on PB and GC cell fate dynamics, by combining combinatorial genetic knockouts with temporally resolved, antigen-specific assays both *in vitro* and *in vivo*. We first tested the hypothesis that NFATC2 and IRF8 antagonize the PB fate by differentiating murine B cells that lack IRF8, NFATC2 or both TFs. Indeed, consistent with our hypothesis, loss of either NFATC2 or IRF8 significantly increased the fraction of cells differentiating into PBs, and combined loss of both TFs produced a larger increase than either individual perturbation (**Figs. 6B and 6C**). Antigen-specific ELISPOT analyses demonstrated an increased frequency of antibody-secreting cells following either individual perturbation, with the largest increase observed following combined loss (**Figs. 6D and 6E**). Consistent with enhanced PB differentiation, *Prdm1* transcript abundance and BLIMP1 protein expression were increased in the perturbed cells, with the strongest effects in the combined knockout (**Figs. 6F and 6G**). Together, these results indicate that both NFATC2 and IRF8 restrain the PB differentiation program and that their combined loss has a greater phenotypic consequence than either single loss. The magnitude of the combined effect was consistent with additive rather than supra-additive contributions, and we therefore describe the two factors as acting cooperatively and non-redundantly rather than synergistically.

We next tested the cooperative action of NFATC2 and IRF8 in GC fating during an in vivo B cell response to immunization with NP-KLH. To do so, we performed competitive adoptive-transfer experiments involving CD45.2 cells that were WT, *Nfatc2*-deficient, *Irf8*-deficient, or *Nfatc2, Irf8*-deficient along with WT CD45.1 cells used as an internal reference population. In response to NP-KLH immunization, loss of either *Nfatc2* or *Irf8* reduced GC B cell proportions and increased PB generation, with the combined knockout producing the most pronounced shift toward the PB state (**Figs. 6H-J**). These results demonstrate the cooperative interplay of NFATC2 and IRF8 in regulating the GC-PB fate bifurcation.

Importantly the altered cell fate distributions caused by the single or combined TF perturbations were accompanied by corresponding changes in the antigen-specific humoral responses. Combined *Nfatc2* and *Irf8* deficiency increased the early antigenspecific IgM response at day 7 (**Fig. 6K**), consistent with enhanced extrafollicular PB differentiation. Conversely, class-switched antigen-specific IgG responses were reduced at day 21, together with a reduction in IgG antibody-secreting cells (**Figs. 6L and 6M**), consistent with a diminished GC response. Finally, the combined loss of *Nfatc2* and *Irf8* resulted in fewer long-lived bone marrow plasma cells that are generated from precursors emanating from germinal centers (**Fig. 6N**). Thus, genetic perturbation *in vitro* and *in vivo* functionally validates a novel role for NFATC2 as a regulator of the GCPB fate bifurcation and supports its cooperative action with IRF8 (**Fig. 6O**). In the updated resulting sub network, NFATC2 and IRF8 constitute a coherent feed-forward loop downstream of BCR signaling that opposes the incoherent IRF4-PRDM1 loop, a topology expected to sharpen the threshold at which activated B cells commit to the PB fate rather than simply shifting the balance between two mutually antagonistic modules^29,30^. These findings thus provide experimental validation of a regulatory relationship predicted by the integration of TF perturbation-derived LFs with inferred GRNs.

## Discussion

Here, we demonstrate the utility of FOCAL, a generalizable framework for prioritizing regulatory subnetworks within state-specific or dynamic GRNs by coupling them to an interpretable ML approach, and of the FOCAL-GIFs that it generates. The fundamental advance of this framework lies in the coupling of complementary lines of inference with distinct assumptions. The ML model, SLIDE, has formal statistical properties regarding identifiability, inference and FDR control under its stated assumptions, and operates solely on the transcriptomic data to identify LFs that discriminate cellular states. SLIDE’s statistical properties have previously enabled discovery of several novel cellular signaling and humoral mechanisms across contexts^14,17-20^. Here, we leverage SLIDE in a regulatory context to simultaneously prioritize both TFs and TF-target linkages.

SLIDE does not utilize any biological priors but instead identifies multigene transcriptional programs associated with alternative cellular states. By contrast, state-specific and dynamic GRNs infer regulatory linkages defining discrete or continuously evolving cellular states using both transcriptomic and chromatin accessibility data and biological priors including information regarding TF motifs and binding. However, GRNs assembled in this manner cannot independently evaluate the regulatory relevance of their own linkages using topology-derived metrics without introducing circularity. FOCAL couples these complementary approaches by intersecting sequence-anchored, mechanistically constrained GRNs with outcome-supervised multigene latent factors. This integration prioritizes regulatory subnetworks that neither approach identifies on its own and mitigates the topological circularity that limits conventional GRN prioritization.

SCENIC+ also uses cell state information to prioritize eRegulons but does through conventional univariate tests. These tests do not provide the same statistical guarantees as SLIDE^14^. FOCAL also differs from approaches based principally on network-topological metrics. Measures such as degree and centrality can effectively identify influential TF nodes, but highly connected TFs may have hundreds of candidate targets, and topology alone provides limited resolution for determining which of those linkages are most relevant to a specific phenotypic contrast. Outcome-associated LFs provide an additional criterion for refining such broad regulons to a restricted subset of TF-target relationships.

Importantly, FOCAL relies on interpretable latent factors. Unlike black-box machine learning approaches where predictive features are obscured within hidden layers, the latent factors generated by SLIDE are statistically identifiable and map directly to coherent transcriptional programs with strict false-discovery-rate control. Consequently, the intersection of these two modeling frameworks yields biologically credible hypotheses that are supported simultaneously by physical priors and outcome-driven covariance. Furthermore, the coupled integration enables the analytical framework to prioritize TF nodes as well as regulatory TF-gene linkages. While topological metrics such as centrality effectively highlight a transcription factor’s global network influence, they cannot specify which of a TF’s hundreds of candidate targets affect a given cellular state or phenotype or when along a differentiation trajectory those linkages become prominent. By evaluating TF-force at the edge level, the FOCAL-GIFs isolate the specific TF-target connections that are predicted to orchestrate cell state transitions.

Applying FOCAL to state-specific GRNs across B and T cell trajectories in the immune system, we were able to prioritize compact regulatory subnetworks whose linkages were more strongly associated with the state contrast than those recovered by alternate approaches. Importantly, our prioritization was successful both at the level of the TFs themselves and the corresponding TF-gene links, across episodes of dynamic TF activity. These prioritized linkages bridge the conceptual gap between stable cellular states and their transition dynamics. State-specific GRNs excel at defining stable architectures but frequently miss transient regulators driving fate bifurcations. However, because the transcriptional signatures of established cellular states retain the legacy of the regulatory events that formed them, outcome-associated latent factors implicitly capture dynamic information. By projecting these stable-state latent factors onto temporally resolved dynamic GRNs, we can retrospectively map fate-associated transcriptional features to the specific, transient regulatory episodes that preceded them.

The predictive capacity of FOCAL was further enhanced by learning directly from TF perturbations. By deriving latent factors from perturbation models, such as the IRF4 and PRDM1 knockouts, and projecting them onto unperturbed progenitor populations, we were able to analyze transcriptional predispositions to alternative cell fates before overt lineage divergence. This enabled the analysis of initiation of dynamic patterning of alternate cellular fates, PB versus GC, within an apparently equivalent group of activated B cells.

The utility and power of FOCAL was demonstrated by uncovering an unsuspected cooperative interplay between the TFs NFATC2 and IRF8, in restraining plasmablast generation and promoting instead the GC fate of antigen activated B cells. Combined deletion of these genes promoted the generation of plasmablasts *in vitro* and profoundly skewed B cell differentiation *in vivo*, during antigen-specific immune responses. The consequence of the combined perturbation in relation to the individual gene deletions was a more pronounced extrafollicular PB response while diminishing the generation of class-switched GC B cells and long-lived bone marrow plasma cells. Crucially, this biological discovery was not made possible by TF-focused GRN centrality. Rather, it emerged from the directed coupling of mechanistically constrained GRNs with outcome-supervised, multigene latent factors.

The NFATC2–IRF8 module extends a regulatory logic for the GC/PB fate bifurcation that has been largely based on pairs of mutually antagonistic transcription factors. Our earlier analyses established that IRF4 and IRF8 antagonize one another and that graded IRF4 concentrations partition activated B cells between the GC and PB trajectories, with high IRF4 levels inducing PRDM1 and the plasmablast program. Work by others has framed the fate decision in complementary terms focusing on antagonism between BCL6 and PRDM1.

What has remained unclear is how signaling input at the time of antigen encounter is coupled to this TF sub-circuit. NFATC2 is a particularly attractive candidate for that role. As a calcium- and calcineurin-regulated factor, its nuclear accumulation is controlled by BCR signal strength and duration, both of which contribute to determining whether an activated B cell adopts an extrafollicular or a GC fate. Notably, NFATC2 was prioritized from latent factors learned from the IRF4 and PRDM1 perturbations rather than simply from its transcript abundance, illustrating the value of target-based inferences for TFs regulated post-transcriptionally. Thus, NFATC2 appears to constitute a B cell receptor signaling input which in conjunction with IRF8 opposes the IRF4-PRDM1 module and its action raises the threshold for plasmablast commitment and favors GC responses.

Several limitations of our framework are worth noting. First, FOCAL prioritizes linkages that are initially specified within a GRN. Regulatory edges that are absent from the assembled GRN cannot be assigned within the latent factors, so the framework improves precision rather than coverage. Second, although the latent factors are selected without reference to network topology or sequence priors, both they and the GRN derive in part from the same transcript covariance structure, so the two evidence layers are complementary rather than strictly orthogonal. Third, the correspondence between LF-prioritized TFs and high perturbation scores is a convergence between two model-derived evidence layers and does not by itself constitute experimental validation. Fourth, the assignment of prioritized linkages to phases is correlative and rests on pseudo-temporal ordering. Finally, the fate predisposition of individual activated B cells is inferred transcriptionally. Clonal barcoding or lineage tracing will be required to test if activated B cells predicted to be predisposed for a particular fate, do so *in vitro* or *in vivo*.

Because FOCAL couples an outcome-supervised evidence layer to an independently inferred interaction network, the framework is not intrinsically restricted to transcriptional regulation. Given suitable paired measurements and a phenotypic contrast, the same coupling could in principle be applied to protein-protein interaction or metabolic networks, generating the corresponding latent-factor-prioritized networks. The FOCALGIFs described here represent one such product, and the generality of the underlying coupling remains to be established in these other settings.

While significant conceptual strides have been made in learning robust embeddings from genomic, transcriptomic and proteomic data, most of these embeddings are either uninterpretable or at best can be weakly linked in a post-hoc fashion to meaningful biological inferences. Our approach, on the other hand, provides interpretable latent representations of regulatory programs. Its generalizability stems from the coupling of two frameworks that rest on distinct forms of evidence, and the corresponding ability to apply these interpretable representations across cell types and states. This is a fundamental departure from deep learning frameworks that attempt to learn an “average” embedding that is useful for many downstream tasks. The ability to resolve robust, context-specific regulatory linkages through complementary evidence layers provides a crucial foundation for higher-order biological modeling and therapeutic engineering of cells. As the field advances toward constructing unified, multi-scale models of complex cellular differentiation trajectories, densely connected correlative networks will not be sufficient. Frameworks capable of distilling large-scale multi-omic data into causal, temporally resolved, and independently validated regulatory subcircuits will be essential for precisely controlling cellular states in health and disease.

## Methods

### FOCAL framework overview

The FOCAL workflow consists of: (1) inferring latent factors (LFs) discriminating cell states using interpretable machine learning modeling; (2) constructing cell-state specific and cellular transition resolved dynamic GRNs, using mechanistic models; (3) enriching for highly-specific dynamic TF activity related to phenotype, and their downstream regulatory linkages; (4) locating the enriched TF-target linkages along the pseudotime trajectory of cellular transitions; (6) extending this framework to lineage defining TF-perturbation derived LFs, for identifying regulatory-network subcomponents associated with cell-fate. We developed FOCAL in Python, with an open-source codebase available on GitHub (https://github.com/jishnu-lab/FOCAL). Analysis notebooks (Python and R) for reproducing the results in this manuscript are at https://github.com/jishnulab/FOCAL_notebooks. Additionally, we provide a user-friendly application (https://pittcsi.shinyapps.io/focal/) for interactive exploration of FOCAL outputs in B-cell differentiation and T-cell exhaustion contexts.

### Inferring interpretable latent factors

SLIDE was used to identify gene programs associated with pre-defined cell states from scRNAseq data. Genes with zero unique molecular identifier counts across all cells were removed, together with mitochondrial and ribosomal genes. Genes with sparsity of counts above a threshold (zero expression in > 30 percent of the cells) across cells were dropped. We further applied SLIDE to single-cell TF perturb-seq datasets to identify perturbation-associated gene programs used to assess predisposition toward alternative cell fates. Model parameters are on Github.

Biological annotations were assigned to the LFs with an LLM-based gene-set interpretation tool (Supplementary Note 1).

### Mechanistically modeling state-specific and dynamic GRNs

#### State-specific static GRN inference

For the B-cell dataset, the base GRN was assembled from MIRA in-silico deletion scores and Cicero co-accessibility links and fitted with CellOracle as previously described^15^. For the T-cell dataset Cell Oracle’s murine base GRN was used due to the absence of ATAC-seq profiles.

Within each cellular transition context (B or T), the two cell-state specific GRNs were merged into a fusion GRN. Edges at or above the 90th percentile of each fusion network’s weight distribution were labeled strong, and the remainder weak. Only strong edges were chosen from the cell state specific GRN’s to create a fusion GRN. In cases where the strong edges were present in both state subtypes unique edge was chosen by retaining the one with higher absolute weight, and strong edges unique to a subtype were carried over unchanged.

#### Pseudo-bulking across trajectory to construct overlapping windows

B-cell differentiation trajectories were inferred with STREAM as previously described (Supplementary Figs. 1 and 2)^15^. For the T-cell system, STREAM was applied to nontargeted control cells from the RBPJ-KO Perturb-seq dataset without imputation. Cells and features were filtered on raw counts, then normalized and scaled, and 370 variable genes were selected by fitting a loess curve (fraction 0.01, 80th percentile) to the per-feature mean–standard deviation relationship. Elastic principal graphs were fitted for both systems with epg_alpha = 0.02, epg_mu = 0.07, epg_lambda = 0.02 and epg_trimmingradius = 2, seeded from 10 clusters, yielding branches on the trajectories and a pseudotime value per cell (Supplementary Fig. 3). Key exhaustion defining TFs expression on the trajectory are displayed (Supplementary Fig. 4)

The fitted flat tree was exported for dynamic network inference. A branch is named by the pair of nodes it connects, and all quantities on a branch (*A,B*) are expressed on the distance-from-nodeaxis, so that branches sharing an origin node (in the B-cell system, the plasmablast branch (0,2) and the germinal-centre branch (0,3), which share the pre-bifurcation portion) are directly comparable. Quantities defined on different origin nodes are never mixed. Cells ordered along the trajectory were tiled by overlapping moving windows (194 windows for the B-cell system), and each window was treated as an independent multi-omic pseudobulk from which one transient-context-specific network was inferred.

For each window, a binary TF → target prior mask was derived from the window’s own ATAC fragments when a matched scATAC-seq profile was available, and otherwise from CellOracle’s ATAC-seq–derived base GRNs (Supplementary Note 2). This mask constrains which edges the network model may fit.

#### Pseudotime-resolved dynamic GRN reconstruction

Dynamic GRNs were reconstructed with adapting a stochastic differential equation (SDE) based framework from Dictys [cite]. A window’s expression matrix and its binding prior mask derived for both joint multi-omic or scRNA-alone datasets, were the inputs to the SDE solver. Networks were reconstructed with distributed GPU compute jobs, with parallelization scripts provided. Reconstruction yields the raw direct edge coefficients per-window, and the normalized *total-effect* network, in which regulation propagating indirectly through the network is accumulated onto each TF–target pair. Unless stated otherwise, all dynamic analyses below use the normalized total-effect weights, so that a transcription factor’s influence on a gene includes the paths that run through its other targets.

#### Smoothed regulatory curves along pseudotime

FOCAL’s temporal module reads the resulting dynamic-network object and resolves it onto a continuous pseudotime axis before any downstream quantity is computed. For a branch delimited by a pair of trajectory endpoints, *M* equi-spaced pseudotime points *t*_1_ <… < *t*_*M*_ are sampled along that branch, and each window *w* = 1,…, *W*, where *W* is the total number of windows used, is located at the pseudotime *τ* _*w*_ of its centroid. A per-window quantity *q*_*w*_ (an expression value or an edge weight) is carried onto the sampled points as a kernel-weighted average over windows,

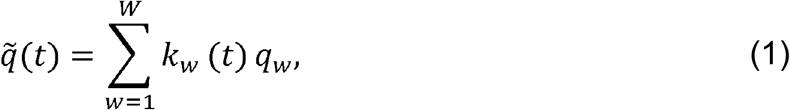

where the kernel weight of window w at pseudotime *t* is a Gaussian kernel of bandwidth *d* in trajectory distance, normalized so that the weights sum to one over the *W* windows at every sampled point,

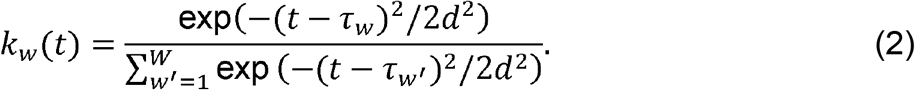

Four families of curves are built accounting for exclusion of any NaN values. Expression curves are the smoothed log CPM of every gene *x*_*g*_ (*t*), whose TF subset *x*_TF_ (*t*) supplies the regulator abundance. Regulation curves are each TF’s out-degree in the binarized smoothed network. Third, beta curves are the signed smoothed edge weight *β* _TF→target_ (*t*) of a queried TF–target link, taken from the normalized total-effect network unless stated otherwise. Lastly, we define a fourth metric called the force curves.

Beta curves are the fitted coefficient *β* _TF→target_ (*t*) measuring how strongly a TF can act on a target gene at pseudotime, but not whether the factor is actually present; conversely, TF expression alone says nothing about which targets a factor engages. We therefore define the *dynamic regulatory force* exerted by a TF on a target as a signed combination of the two,

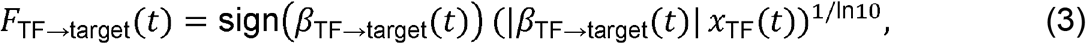

where *x*_TF_ (*t*) is the smoothed log_2_ CPM expression of the TF (Supplementary Fig. 5). The force is the product of regulatory strength and regulator abundance raised to the power 1/ln10 ≈ 0.434; the implementation evaluates it as exp [log_10_ |(*β*| + *ε*) + log_l0_ (x_TF_ + *ε*) with *ε* = 10^−10^ guarding the logarithms at zero, which is the same quantity up to. The force is therefore zero when either TF or gene is absent, grows with both, inherits its sign from *β*—positive force denotes activation and negative force repression—and the sub-linear exponent keeps edges whose coefficients span several orders of magnitude on a comparable scale. Evaluated over the sampled points of a branch, F_TF→target_ (*t*) forms the *force wave* of that link. Two summaries of a force wave are used below: its mean over an episode, 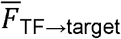, which ranks edges when an episodic GRN is assembled, and its absolute maximum along a branch, max_*t*_|F_TF→target_ (*t*)|, which is the per-link statistic used for phase assignment and for validation.

Force waves are inspected in two complementary ways. For a single link, the force function is drawn as a surface over regulator abundance *x*_TF_ and coefficient magnitude |*β*|—one sheet for activation and one for repression—and the link’s own wave is traced on that surface, so that the path a link takes through the (abundance, coefficient) plane and the force it generates are read off together. For sets of links, the waves are stacked into a force heatmap over pseudotime on a symmetric diverging scale, so that activating and repressing programs are directly comparable; rows (and, where informative, columns) can be ordered by hierarchical clustering of the waves themselves (Ward linkage on Euclidean distance), which groups links that rise and fall together regardless of which TF drives them.

Window pseudotimes and sampled-point pseudotimes are placed on a common scale with AlignTimeScales, so that window-level quantities (cell-state composition and TF binding, described below) and point-level quantities (expression, *β*, force) can be compared on the same axis. Unless stated otherwise, curves were sampled at *M* = 40 points with *d* = 0.001 in the B-cell system and at *M* = 40 points with *d* = 0.002 in the T-cell system; force waves in the B-cell system were resolved at a denser *M* =100 points with *d* = 0.0005.

### Construction of GRNs across episodes of TF activity during cellular state transitions

A single GRN fitted over an entire transition trajectory average over regulatory programs that are active at different times. We therefore partition each branch into consecutive *episodes* and reconstruct a GRN within each one. An episode is a contiguous block of sampled pseudotime points; with *M* = 40 points and 5 points per episode this gives 8 episodes per branch, and 4 episodes per branch were carried forward for the plasmablast (PB) and germinal center (GC) analyses (Supplementary Fig. 6). The linear T-cell trajectory was sampled at *M* = 24 points with 4 points per episode, giving 6 consecutive episodes (Supplementary Fig. 7).

Within an episode the smoothed weighted network is sliced to that episode’s points and unrolled into a TF × target edge table with one column per point. Edges that are identically zero across the episode (edges with no ATAC-seq support in the base GRN) are dropped, and regulators whose gene symbols begin with ZNF or ZBTB are excluded, as these large paralogous families lack the motif resolution to be assigned confidently.

Each retained edge is then tested for *temporal invariance* across the episode. For edge TF→target let (*β*_1_,…, *β*_*m*_) be its non-zero coefficients over the episode’s points. The edge is retained when at least 3 of the episode’s points carry a non-zero coefficient; a one-sample-test of those coefficients against zero is significant (p < 0.05 at the filtering stage, tightened to p < 0.001 for the episodic GRN); and the direction of regulation is invariant, i.e. all non-zero coefficients share the same sign, so that a factor does not switch between activating and repressing its target within a single episode.

An edge that passes all three criteria represents a regulatory relationship that is both statistically supported and directionally stable for the duration of the episode. For every retained edge the force curve F_TF→target_ (*t*) is computed from that edge’s coefficients and the TF’s expression over the same points, and its mean force 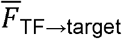 is taken as the edge weight of the episode. The *episodic GRN* is the set of edges in the top 2% of 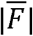 (i.e. 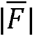 at or above the 98th percentile). Where activating and repressing programs were compared separately, the top 2% of positive and the bottom 0.5% of negative mean forces were selected instead, so that repressive edges—which are far fewer—are not lost to a single two-sided threshold. Episodes are reconstructed independently and in parallel, one process per episode, so that no episode’s filtering influences another’s.

### Transient regulatory activity analysis

To summarize how a TF’s activity is distributed over an entire branch rather than within one episode, each smoothed curve *y*(*t*) (expression or regulation, i.e. log out-degree) sampled at pseudotimes *t*_1_ <… < *t*_*M*_ is reduced to four characteristics on a pseudotime axis rescaled to [0,1]. Writing AUC (*y*) for the trapezoidal integral of over that axis, and

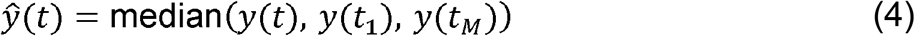

for the curve clipped to the band spanned by its endpoints, the characteristics are the terminal log fold change Δ_term M_= y(*t*_*M*_) − (*t*_1_), the transient log fold change Δ_trans_ = AUC (*y* − *ŷ*), the switching time 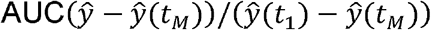, which locates the pseudotime at which the main transition occurs, and AUC (*y*) itself. The terminal term captures net change between the ends of the branch; the transient term captures excursions away from the endpoint band and is by construction insensitive to net change.

Both terms are - scored across all curves, and each TF is assigned the wave pattern of whichever term dominates in absolute value: *up* and *down* when |*z* (Δ_term_) | ≥ |*z* (Δ_trans_)| (sign of Δ_term_1 deciding between them), and *transiently up* (a bell-shaped wave) and *transiently down* (a U-shaped wave) otherwise. TFs are additionally ranked within each pattern by the absolute*z*-score of the dominant term, which is how the top regulators per pattern were selected for display. Transiently active regulators identified this way are exactly those a terminal-fold-change analysis, and any comparison of the two endpoint states, would miss.

### Chromatin binding inference across pseudotime trajectories

TF binding along each branch was quantified directly from the per-window chromatin footprints produced during dynamic-network inference, giving smoothed per-TF binding-score and bound-OCR-count curves on the branch’s pseudotime axis (Supplementary Note 4, Supplementary Fig. 8).

### Cell-state composition along the trajectory

The dynamic network assigns each cell to one or more pseudotime windows. Combining this soft cell-to-window assignment with the cell-state labels, we tabulate how many cells of each state fall in every window, giving a state × window count table, and place each window on the branch’s pseudotime axis with AlignTimeScales. The progenitor states shared by all branches (ActB-1, earlyActB) are dropped, as they contribute to every lineage alike and would mask lineage-specific structure. Restricting the table to the windows of one branch gives that branch’s composition trajectory, which we display both as per-state count curves over pseudotime and as stacked bars of the average composition over equally spaced bins of windows.

Each state’s count trajectory is then reduced to its extrema: maxima are located by prominence-based peak finding on the trajectory and minima by the same procedure on its negation (prominence of 10 cells, minimum separation of 3 windows). A state’s maximum marks its peak abundance along the branch and the first minimum after it marks its collapse; the pseudotimes at which states collapse are the cell-state switches that delimit the regulatory phases defined next.

### Prioritizing state-specific TFs and linkages most associated to phenotype

To identify TFs with regulatory programs enriched in cell subtype discrimination specific regulons, we performed enrichment analyses using previously constructed fusion GRNs alongside regulons inferred from SLIDE. By overlapping GRN topology with both standalone and interacting regulons, we assembled a comprehensive set of TF–target gene relationships. These interactions were then evaluated through two levels of enrichment analyses. The order-1 (single-TF) enrichment analysis, in which for each regulon, TFs either directly included in the regulon or connected as first-degree neighbors (based on GRN topology) were considered. To assess their regulatory importance, we quantified how many genes each TF regulates strongly within the regulons compared to its global regulatory profile in the fusion GRNs. For each TF, the enrichment score was computed as:

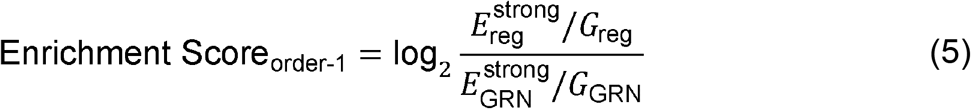

The enrichment score thus quantifies the TF’s overrepresented strong regulatory influence for given regulons. It is reported on a log scale, so that a score of zero denotes a regulon in which the TF’s strong targets occur at exactly the frequency expected from the fusion GRN, positive scores denote over-representation, and each unit is a doubling of the observed-to-expected ratio. A hypergeometric test was used to determine statistical significance, and transcription factors with significantly enriched regulons were identified as candidate key regulators.

### Enriching for dynamic TF activity in episodic GRNs

The trajectory branches for both datasets were broken into episodes as described above; the B-cell system bifurcates into the PB and GC branches, whereas the T-cell trajectory is linear, so a single ordered series of episodes spans it. Given an episodic GRN and a latent factor (its gene set), we ask which transcription factors regulate that program disproportionately within that episode.

Let *N* be the number of distinct target genes in the episodic GRN, *K* the number of program genes that are active as targets in that episode, *n*_TF_ the number of targets of the TF in the episodic GRN and *k*_TF_ how many of those belong to the program. Enrichment is scored as the fold over-representation

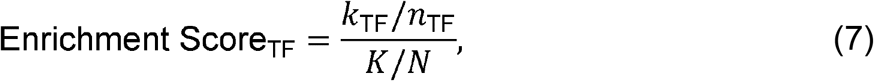

and its significance is assessed with a one-sided hypergeometric test,

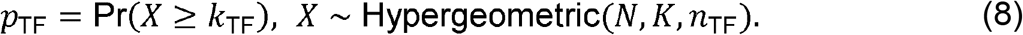

TFs are ranked by enrichment score, those with a score of zero are discarded, and significant regulators are retained (*p* < 0.05). For the episodic enrichment dot plots a stricter, lineage-specific cutoff on the raw p-value was applied—*p* ≤ 0.003 on the PB branch and *p* ≤ 0.01 on the GC branch—and a factor is displayed only if it clears that cutoff in at least one episode. Reported *p*-values are raw hypergeometric tail probabilities; no multiple-testing correction is applied, and the cutoffs above should be read as display thresholds rather than as controlled error rates.

Note that the episodic enrichment score is a linear fold over-representation, unlike the log fold-enrichment used for the state-specific order-1 and order-2 analyses above; the two effect sizes are on different scales and are not pooled. In the episodic enrichment dot plots the two channels encode the two statistics separately: dot colour encodes the enrichment score and dot area encodes significance, ramping linearly in − log_10_*p* between the display threshold and a cap at *p* = 10^−6^, with non-significant entries drawn at a fixed minimal size. Where a dot plot spans a wide range of enrichment scores, colour is optionally rendered on a log_2_ scale for legibility (with a constant offset added beforehand if any score is non-positive, as annotated in the colour-bar label); this is a display transform only and does not change the reported scores or their significance. For each enriched TF the analysis also returns the program genes it targets in that episode together with their mean forces, and its remaining downstream targets, so that the sign and strength of each regulatory relationship can be inspected alongside the enrichment statistic. Running this across consecutive episodes yields the episodic enrichment pattern of a TF—the sequence of episodes in which it acts on the program—which distinguishes factors that act throughout the transition from those that act only within a restricted window of it.

Because episodes are reconstructed independently, an individual TF–target edge can also be followed across them. For a chosen set of TFs and targets, each edge’s mean force is collected from every episodic GRN into an edge × episode matrix, with zero entered wherever the edge did not survive that episode’s filtering; this matrix is what the TF–target episodic heatmaps display. In the episodic enrichment dot plots, a TF is shown for an episode only if it targets at least two program genes and at least two further downstream genes there. Unless an explicit order is imposed, TFs are ordered by the episode in which they peak—the significant episode with the highest enrichment score, falling back to all episodes when none is significant—and, within an episode block, by decreasing peak enrichment score. Alternatively, TFs are ordered by the similarity of the program genes they target, taking the Jaccard index between their target sets as the similarity, 1− Jaccard as the distance, and Ward linkage for the hierarchy. The same TF–gene incidences, counted across episodes, give the TF–gene co-regulation map, which shows which factors converge on the same program genes and in how many episodes they do so.

### SCENIC+ benchmarking of FOCAL enriched TF-centric network components

Direct comparison of transcriptional programs enriched by FOCAL TF-centric regulons was done using SCENIC+. We generate state-specific regulons distinguishing GC and PB states in the B-cell system (Supplementary Fig. 9), and Tex-term and Tex-KLR states in the T-cell system (Supplementary Fig. 10). Our workflow proceeded as follows: SCENIC+ was first applied to single-cell multi-omic data of both systems to infer regulons differentially active between the specified cell states. Regulon activity and specificity were quantified using the *eRegulon* score. We retained only those SCENIC+-derived regulons whose TFs overlapped with significantly enriched TFs identified through our regulon and fusion GRN analyses ensuring comparability across analyses. To compare the specificity and sensitivity of those TF-centric regulons, we compiled the target genes of each transcription factor within the selected SCENIC+ regulons, as well as those defined by the regulon and fusion GRN frameworks. We then calculated the absolute fold changes in gene expression between the two cell state subtypes to assess the differential regulatory impact across methodologies.

### Dictys benchmarking of FocalGIFs enriched TF-Target linkages

We benchmark for FocalGIFs-nominated regulators against Dictys, which reconstructs lineage-resolved dynamic gene regulatory networks, nominated TFs. For each lineage branch (B cell: PlasmaBlast and Germinal Center), we ran Dictys’ draw_discover function in regulation mode, which ranks TFs along shape-based metrics. For each branch, the top 5 TFs by each of the four metrics were retained (ntops = (5,5,5,5)), and the union across all four branches and both cell types was taken as the Dictys-prioritized TF set (71 unique TFs, Supplementary Fig. 11).

FocalGIFs-nominated regulators were benchmarked against this set using TF-level in silico perturbation scores generated with CellOracle, in which simulated knockout of each TF yields an overall perturbation magnitude score reflecting its predicted transcriptomic impact. FocalGIFs TFs were partitioned into (i) dynamic TFs, defined as those temporally enriched across the corresponding branches by episodic-enrichment analysis (p ≤ 0.05), and (ii) state-specific TFs, defined by latent factor (LF) enrichment for each terminal state. For each comparison, one-sided Mann-Whitney U tests (alternative = “greater”) were used to test whether FocalGIFs TFs had higher perturbation magnitude than (a) the Dictys-prioritized TF set (excluding TFs shared between the two lists) and (b) all remaining background TFs; effect sizes were reported as Cliff’s delta. The same comparison (dynamic vs. state-specific FocalGIFs TFs) was additionally performed to assess whether state-specific and dynamic programs carried distinct perturbation signatures. Analyses were performed independently for the B cell and Tonsil datasets.

### Deriving phases of gene-regulation governing state switching of cells

Episodes divide a branch into equal blocks of pseudotime; *phases* instead divide it at the points where the cellular composition of the trajectory changes, so that each phase corresponds to an interval between two cell-state switches.

Phase boundaries are derived from the cell-state composition trajectories described above, restricted to post-bifurcation windows so that the boundaries are lineagespecific. For a state *c*, its *termination pseudotime* is the first pseudotime after the state’s peak abundance at which its count falls to a fraction τ of that peak (*τ* = 0.1); alternatively the first local minimum after the peak, located by peak-finding on the negated count trajectory, can be used, falling back to the threshold rule when no such minimum exists. If a state does not collapse within the branch, the branch’s final pseudotime is used, so the rule always returns a usable boundary. In the B-cell system the PB branch is delimited by the terminations of ActB-4 and earlyPB, giving three phases, and the GC branch by the termination of ActB-3, giving two; in general *N* switches define *N*+ 1 phases, and a pseudotime *p* falls in phase *k* when switch_*k*−1_< *p* ≤ switch_*k*_.

Each regulatory link is then assigned to the phase in which its force peaks. Rather than taking the single largest point of a force wave, which is sensitive to the sampling of pseudotime, the peak is estimated by softmax weighting of the wave’s absolute values,

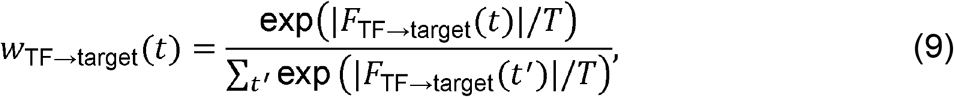

with temperature *T* controlling how peaked the weighting is. The *k* highest-weighted points (*k* = 5) are retained and the link’s peak pseudotime is their weight-normalized mean, 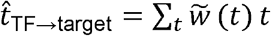, where 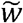 renormalizes the softmax weights over the retained points. (Taking the single top-weighted point, or the unweighted mean or median of the retained points, is also supported.) Binning 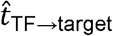 against the phase boundaries assigns each link a phase, and ordering links by 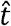 orders them by when their regulation acts. Because the phase is defined as the interval containing the peak, max_*t*_|*F*_TF→target_ (*t*)| is simultaneously the link’s in-phase peak force.

The same phase partition is applied to the chromatin binding curves (Supplementary Note 4). A TF’s *per-phase binding score* is the maximum of its smoothed binding-score curve over the windows falling within that phase—the binding analogue of the absolute maximum force—and TFs are ranked by that score independently within each phase, so that selections are never pooled across phases. This provides an unbiased, per-phase selection of the most strongly binding factors within a category (for example state-specific versus episodic TFs), replacing hand-picked panels. Where two categories are compared, TFs shared between them can be removed from both so that each category is sampled only from the factors unique to it, a size-matched random control is drawn without replacement from the remaining factors scored in that phase, and the two categories are compared within each phase with a two-sided Mann–Whitney *U* test.

### Validation of dynamic TF activity in phases against random regulatory links

To test whether the links prioritized by FOCAL carry more regulatory force than expected by chance, each enriched link was compared against a size-matched null of non-enriched links using the absolute maximum force max_*t*_|F_TF→target_ (*t*)| as the statistic—the same quantity by which links were selected.

Null links were drawn in one of two ways. In the first, a candidate pool of *n*_TF_ × *n*_target_ links is sampled from the network universe after removing every enriched TF and every enriched target (or, in the more permissive variant, only the exact enriched TF–target pairs), the pool is scored, and a random subset matched in size to the enriched set is retained. In the second, links are sampled from edges that are actually present—nonzero in the fitted network—in windows whose cellular composition exceeds a threshold for a given set of states, with all enriched TFs and targets masked out first; force is then scored post hoc. The second null is the stricter of the two, since it holds fixed the fact that an edge exists in the relevant cell states and asks only whether its force is comparable.

Because the B-cell trajectory bifurcates, each link can be scored either on one lineage or across both. In the combined mode each enriched link is scored on the lineage on which its absolute maximum force is larger, and that lineage is recorded alongside the value; the random null is then the union of the per-branch forces of the sampled links, so the null is never given the same cross-branch maximum advantage as the enriched set. Several enriched sets (for example state-specific versus episodic links) can be compared against one shared null, drawn after removing the TFs and targets of all sets and size-matched to the largest set.

The comparison is also carried out within phases. Each enriched link is assigned to a phase of its winning lineage by the softmax peak rule above, random links are scored and phase-binned on the same lineage with identical softmax parameters, and within every (lineage, phase) cell the null is size-matched to the largest enriched set in that cell. This keeps enriched and random links on the same footing at every step: the same network variable, the same force definition, the same peak rule, and the same phase boundaries.

### Calculation of perturbation scores by in silico TF knockout

To quantify the predicted influence of individual transcription factors (TFs) on cell-state transitions, each TF represented in the state-specific GRN was perturbed in silico, one at a time, using CellOracle. For each perturbation, the resulting displacement of individual cells was compared with the differentiation vector field, yielding a per-cell perturbation score. The sign of the score indicates whether the simulated TF knockout shifts a cell along or against the direction of differentiation, whereas the score magnitude reflects the predicted strength of the perturbation. The complete analysis was independently repeated in a human tonsil B-cell dataset. FOCAL-prioritized state-specific and episodic TFs were then compared with the remaining TFs using a onesided Mann–Whitney *U* test.

### Utilization of optimal transport for validating dynamic TF activity

Transcription factors were ranked from optimal-transport couplings between consecutive sampling windows of the B-cell culture, computed with moscot v0.5.0 [cite] on a joint RNA/ATAC embedding of the cells with paired scRNA-seq and scATAC-seq profiles learned with MultiVI [cite] (scvi-tools v1.4.1; default architecture with an 11-dimensional latent space, trained for 200 epochs with a 10% held-out validation split; Supplementary Fig. X). The couplings, the push-forward and pull-back distributions derived from them, and the resulting TF rankings are described in Supplementary Note 5.

### Prediction of perturbation-associated states

We trained L1-regularized logistic regression models, to identify transcriptomic features that distinguish PRDM1- or IRF4-perturbed cells from unperturbed controls. Each trained classifier was subsequently applied to early activated B cells (ABCs) from the multi-omic dataset at days 2 and 4 to estimate, for each cell, the probability of assignment to the corresponding perturbation-associated state. Cells with a predicted probability > 0.5 were classified as perturbation-associated, whereas cells with probabilities ≤0.5 were classified as control-like. The analysis was repeated using only differentially expressed genes between KO and control cells (|log_2_ FC|> 1, adjusted *p* < 0.05). Lasso logistic regression models were trained on this reduced feature set and applied to early ABCs from days 2 and 4 to estimate perturbation-associated probabilities. We next projected SLIDE-derived latent factors from the B-cell perturb-seq data onto day 2 and day 4 ABCs from the single-cell multiome dataset to assess transcriptional predisposition toward GC or PB differentiation. Program scores were generated using the predZ function in SLIDE, quantifying the similarity of each ABC to the corresponding KO-or control-associated state. Using thresholds defined from the perturb-seq models, ABCs were classified according to their inferred transcriptional predisposition. The same framework was also applied to T cells. We enriched for TFs from the episodic GRNs for these perturbation-associated (PRDM1-, IRF4-) gene programs in B-cells and precursor-exhausted T-cell states. (Supplementary Figs. 12 and 13),

Gene set enrichment of the predicted KO-like subsets was assessed with ESCAPE (Supplementary Note 6).

### Mice

C56BL/6J, μMT and *Irf8*^−/−^ mice were obtained from the Jackson Laboratory. *Nfatc2*^*−/−*^ mice were a kind gift of Dr. Anjana Rao, La Jolla Institute for Allergy and Immunology. Mice were housed and bred in specific pathogen–free conditions, in accordance with guidelines of the Institutional Animal Care and Use Committee.

### B cell isolation, cell culture and differentiation assays

Resting FO B cells of the indicated genotypes were isolated using Miltenyi Biotec kit according to the manufacturer’s protocols^29^. B Cells (0.5 × 106 cells/ml) of indicated genotypes were stimulated to differentiate into plasmablasts with F(ab’)2 fragment goat anti-mouse IgM, anti-CD40, IL-2, IL-4 and IL-5 as described earlier^32^. Antibody secreting cells (ASCs) were analyzed by flow cytometry and ELISPOT analyses, respectively

### Adoptive transfer of B cells and NP-KLH immunization

Adoptive transfer experiments were based on a previously described protocol^29^. 1 × 10^7^ resting FO B cells (WT CD45.1^+^: WT, *Irf8*^−/−^, *Nfatc2*^*−/−*^ *or Irf8*^−/−^*Nfatc2*^*−/−*^ CD45.2^+^) and 3 × 10^6^ WT CD3 T cells were co-transferred into 6- to 8-week-old μMT recipients via injection into the tail vein. After 2 weeks, mice were immunized intraperitoneally with NP (29)-KLH (1 mg ml^-1^, Biosearch Technologies) mixed with 50% (vol/vol) alum (Thermo Scientific) and LPS (0.01 mg ml^-1^, Sigma). On day 14 post immunization, naïve, GC B cell and plasmablast populations in the CD45.1^+^ and CD45.2^+^ compartments were analyzed by flow cytometry. Antibody and ELISPOT analyses were performed at D7 and D21 post-immunization as described earlier^32^.

### Flow cytometry analyses

Cells were washed and prepared as single cell suspensions in MACS buffer (PBS plus 1% FBS and 5 mM EDTA), following by blocked with 1% donkey serum and 25μg/ml 2.4G2 (BD) for 15 minutes on ice before adding specific antibodies for cell surface antigens. Cells were typically stained at 4°C for 30-90 minutes. The reagents for flow staining included BV510 anti-B220, FITC anti-GL7, BV421 anti-CXCR4, APC anti-CD138, PerCP anti-CD45.1, AlexaFluor 700 anti-CD45.2, viability Dye eFluor780 from eBioscience, APC anti-CD21, PE-Cy7 anti-CD23.

### Immunoblotting

Proteins in total cellular lysates were resolved using SDS-PAGE and transferred to a PVDF membrane. The blots were probed with □-Blimp1 antibody (C14A4; Cell Signaling Technology), visualized according to supplier protocol and quantitated with an Odyssey Infrared Imaging System.

## Supporting information

Combined supplementary notes and figures

## Acknowledgments

J.D. and H.S. were supported in part by NHGRI U01HG012041. This work was carried out through a Networks Award (U01HG012041) within the NHGRI-funded IGVF Consortium.

## Extended Data Figures

**Extended Figure 1:**
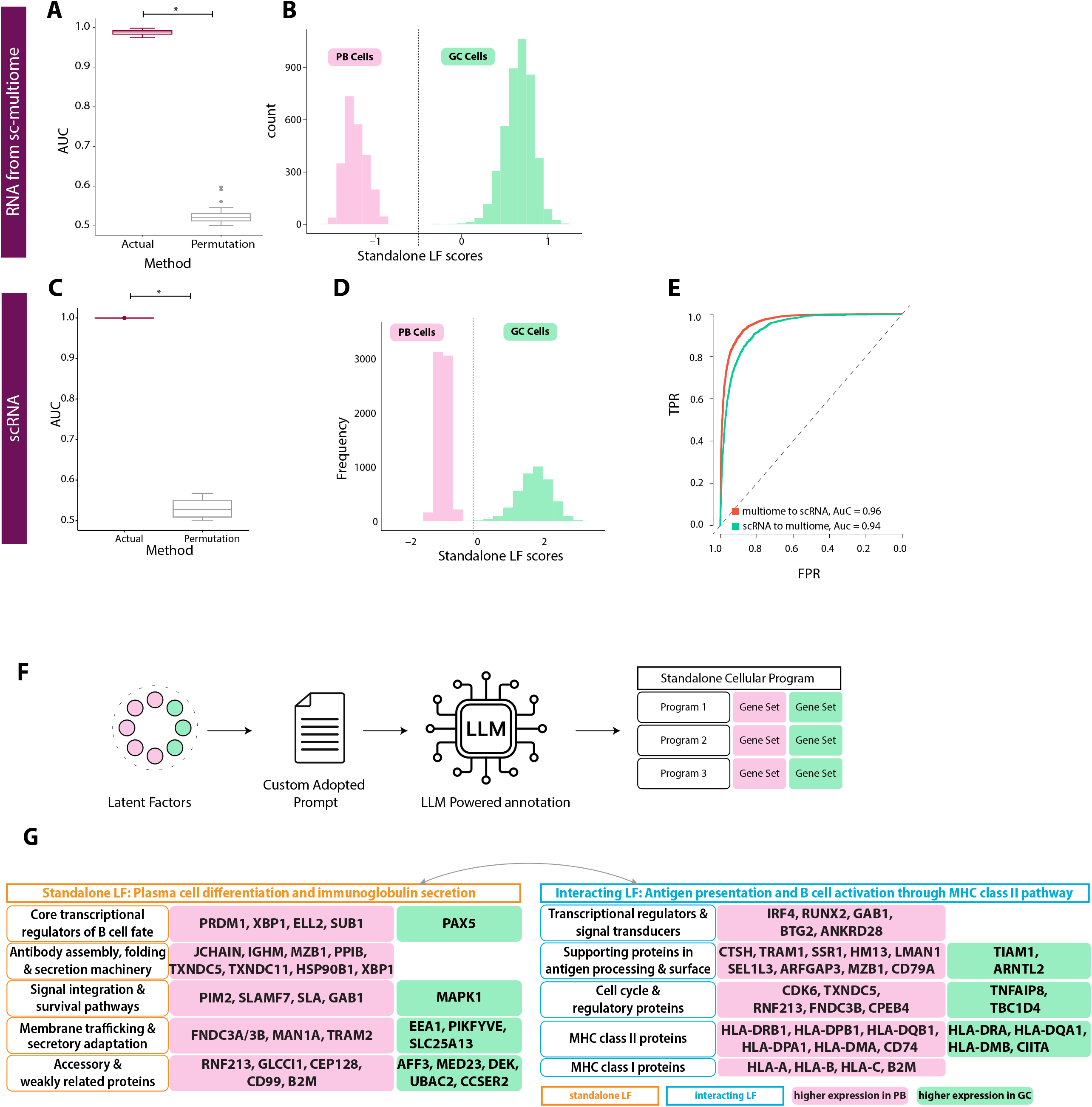
Evaluation of robustness of SLIDE-derived LFs in discriminating GC and PB states and functional annotation of the LFs. **A**. Performance of the SLIDE model built on the scRNA-seq component of the multiome. Model performance evaluated across replicates of k-fold cross-validation and significance evaluated using permutation testing. **B**. Significant standalone LF scores from model in A distinguish PB and GC B cells. Distribution of significant cellular program scores from the model trained on PB and GC B cells, with PB (pink) and GC (green). **C**. Performance of SLIDE model built on an orthogonal scRNA-seq dataset. Model performance evaluated across replicates of k-fold cross-validation and significance evaluated using permutation testing. **D**. Significant standalone LF scores from model in C distinguish PB and GC B cells. Distribution of significant LF scores from the model with pink corresponding to PB cells and green corresponding to GC cells. **E**. Performance of SLIDE models trained on multi-ome scRNA-seq data and tested on scRNA-seq-only data (AUC = 0.96), and vice versa (AUC = 0.94). SLIDE models are trained to distinguish GC and PB cells using transcriptomic data. **F**. Schematic illustrating how the identified LFs are functionally annotated using an LLM-based gene set enrichment analysis. Consensus annotations for each LF is obtained using a pretrained BERT model. **G**. Functional annotation of the LFs distinguishing. PB and GC. Green and pink indicate higher expression in GC and PB, respectively.

**Extended Fig 2:**
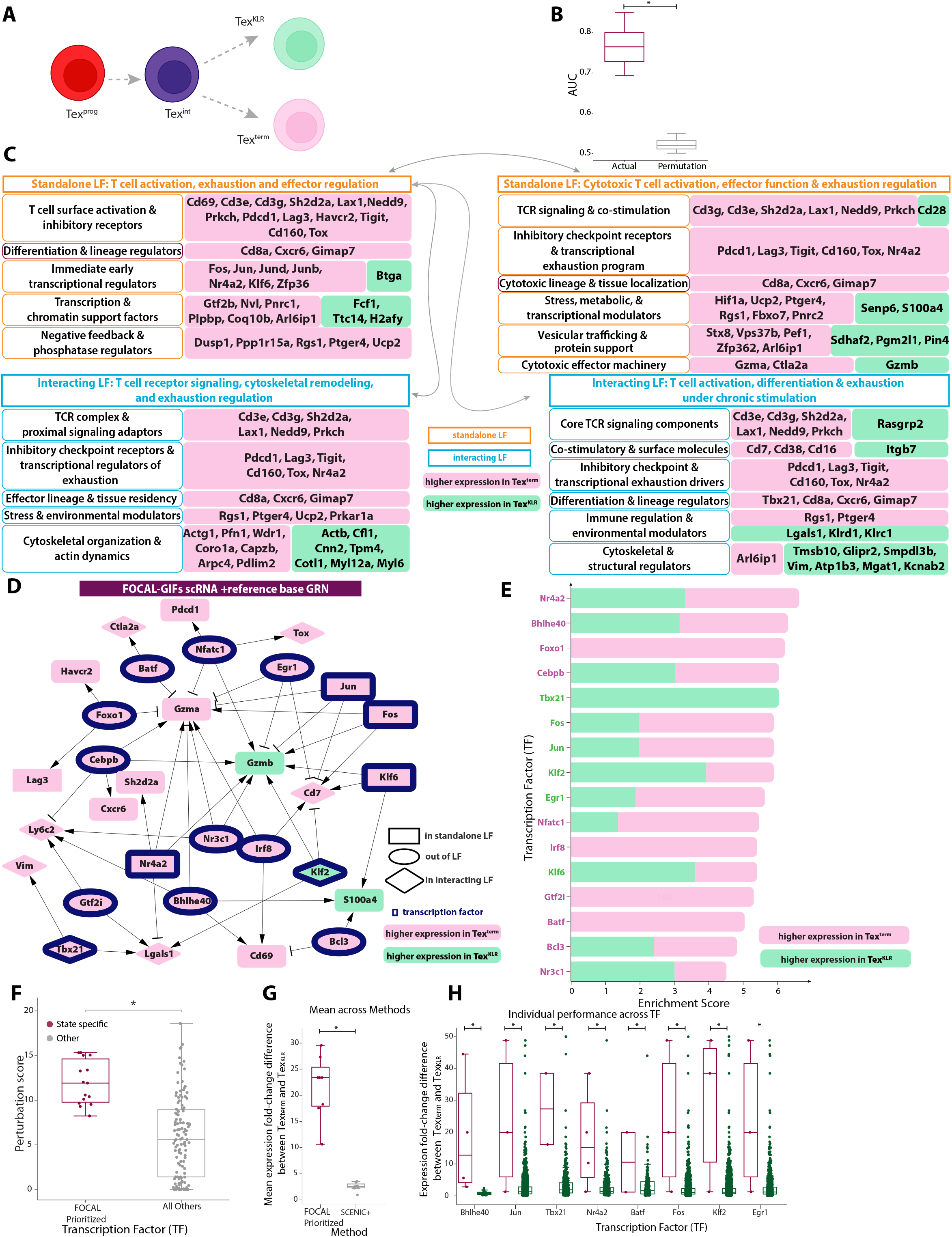
Coupling interpretable ML to state-specific GRNs to prioritize candidate regulatory subnetworks in CD8 T cells. **A**. Schematic of CD8^+^ T cell progression toward exhaustion, diverging into terminally exhausted (Tex-term) and KLR-expressing cytotoxic (Tex-KLR) states. **B**. Performance of SLIDE model built on the scRNA-seq data from Tex-KLR and Texterm cells to distinguish these CD8 T cells subsets. Model performance was evaluated across replicates of k-fold cross-validation and significance evaluated using permutation testing. **C**. Functional annotation of the LFs distinguishing Tex-term and Tex-KLR. Green and pink indicate higher expression in Tex-KLR and Tex-term, respectively. **D**. Coupling SLIDE-derived LFs to state-specific GRNs identifies a prioritized regulatory subnetwork. Colors indicate expression levels in CD8 T cell states (higher expression in Tex-KLR: green, higher expression in Tex-term: pink). TFs are denoted by a blue boundary. TFs/genes in standalone and interacting LFs demonstrated by rectangles and rhombuses respectively. TFs outside LFs are represented using oval shapes. **E**. Enrichment scores for the prioritized TFs from the state-specific GRNs. Colors indicate fraction of downstream genes for that TF with higher expression in the Tex-KLR (green) or Tex-term states (pink). **F**. Perturbation scores for prioritized TFs using in-silico perturbations on state-specific GRNs in Tex-term and Tex-KLR CD8 T cells. **G**. Distribution of mean absolute logFC of gene expression values of downstream target genes for TFs shared between FOCAL and SCENIC+. Each point represents the average logFC (Tex-term:Tex-KLR) value for a TF across all the prioritized TF-gene links. **H**. The full distribution of absolute logFC values for downstream genes of each shared TF as inferred by FOCAL and SCENIC+. Each point represents a TF-gene link-specific logFC (Tex-term:Tex-KLR) value.

**Extended Figure 3:**
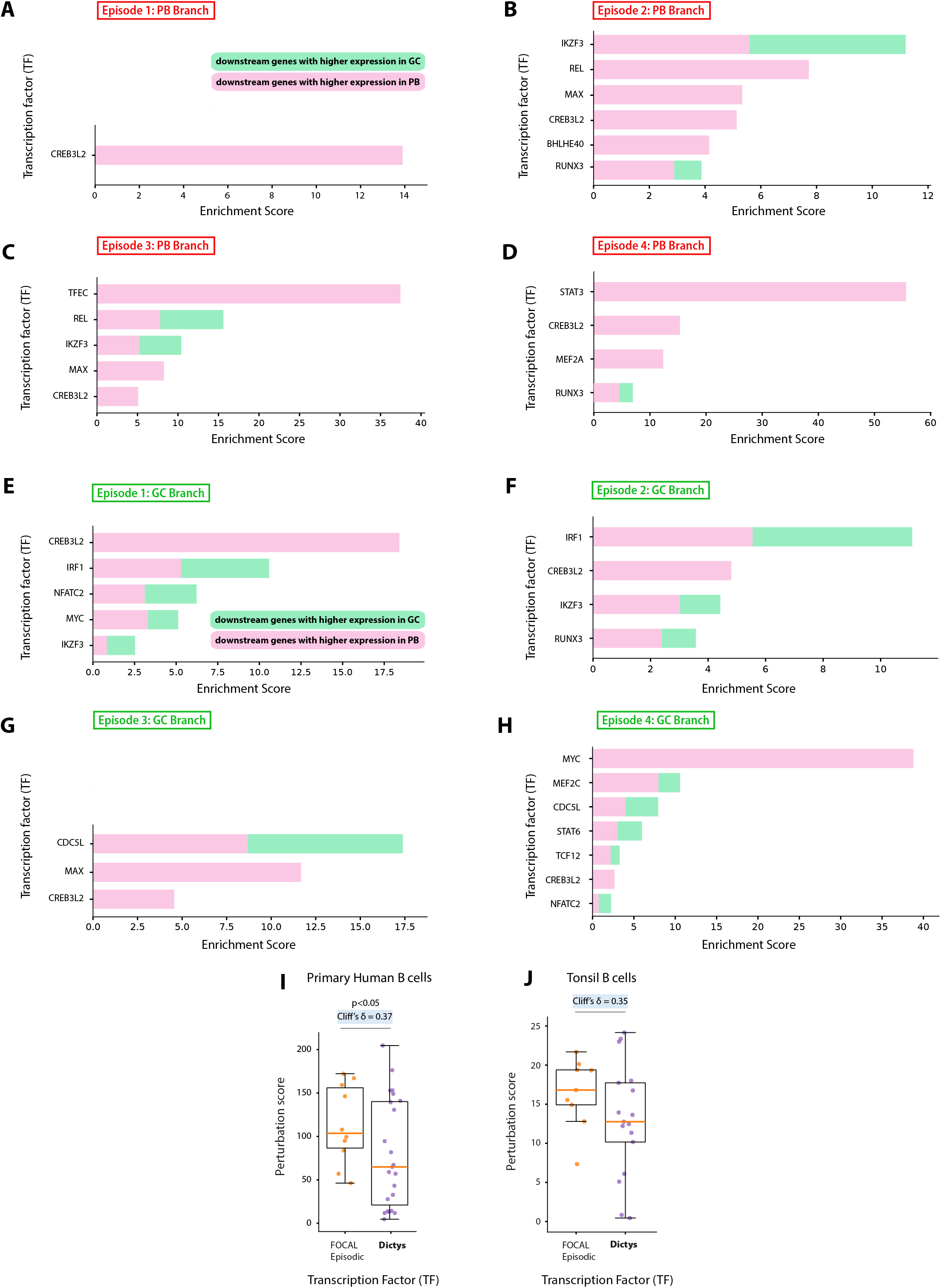
Additional evaluation of TFs prioritized by FOCAL. **A, B, C, D**. Enrichment scores of prioritized TFs per significant episode of activity for the PB branch. Colors indicate fraction of downstream genes for that TF with higher expression in the GC (green) or PB states (pink). **E, F, G**. Enrichment scores of prioritized TFs per significant episode of activity for the GC branch. Colors indicate fraction of downstream genes for that TF with higher expression in the GC (green) or PB states (pink). **H, I**. Benchmarking of FOCAL prioritized TFs with Dictys prioritized TFs using perturbation analysis on single-cell multi-ome B cell and human tonsil atlas data.

**Extended Fig 4:**
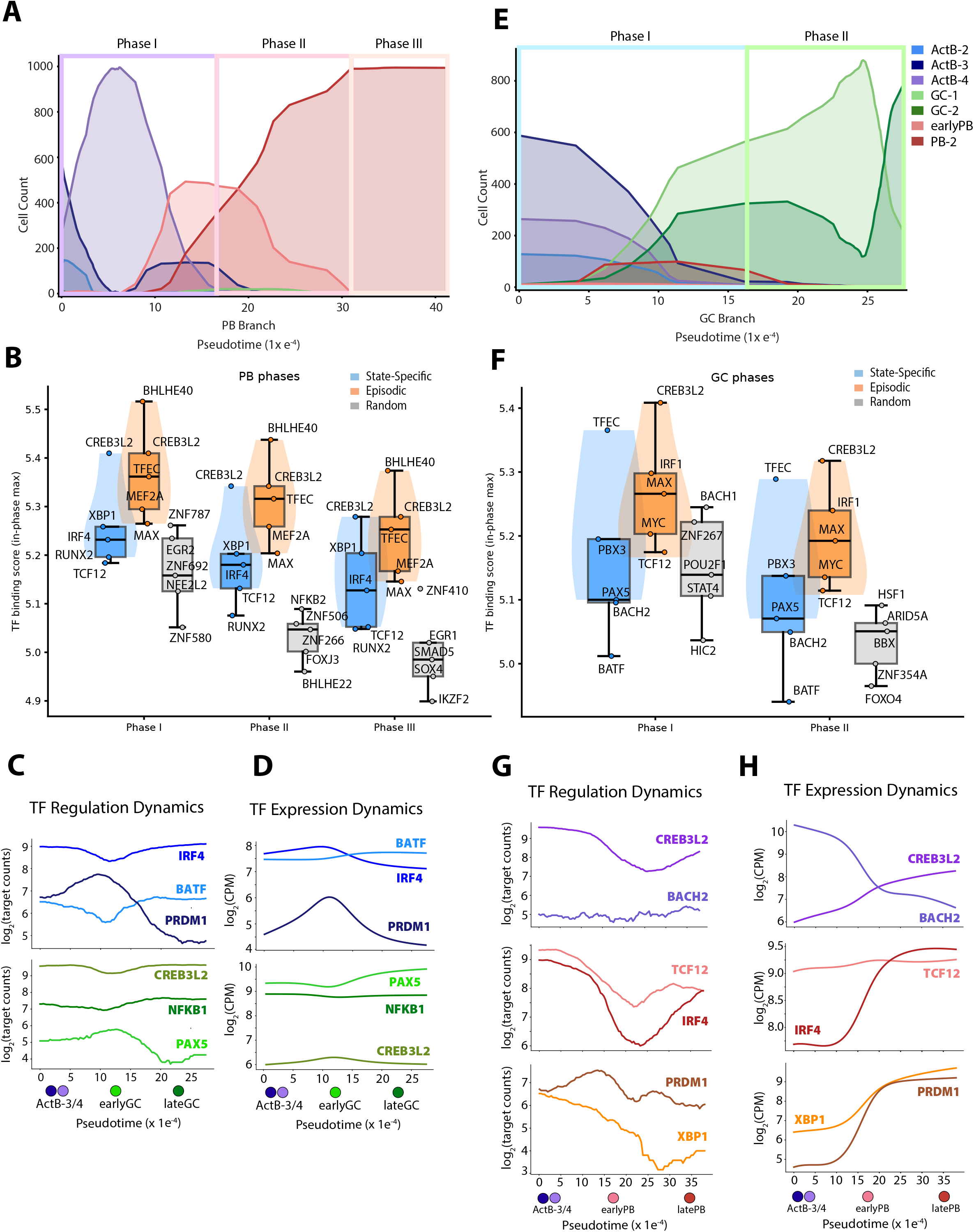
TF binding, regulation and expression dynamics for prioritized TFs across the PB and GC branches. **A**. Phase-specific cell populations for the PB branch. Curves reflect frequencies of different populations of B cell states across the three phases in the PB branch. **B**. TF binding scores for TFs prioritized from the state-specific or dynamic GRN analyses vs random TFs across phases in the PB branch. **C**. Regulation dynamics (quantified using network connectivity over pseudotime) of key prioritized TFs in the PB Branch. **D**. Gene expression dynamics (quantified using expression levels over pseudotime) of key prioritized TFs in the PB Branch. **E**. Phase-specific cell populations for the GC branch. Curves reflect frequencies of different populations of B cell states across the two phases in the PB branch. **F**. TF binding scores for TFs prioritized from the state-specific or dynamic GRN analyses vs random TFs across phases in the GC branch. **G**. Regulation dynamics (quantified using network connectivity over pseudotime) of key prioritized TFs in the GC Branch. **H**. Gene expression dynamics (quantified using expression levels over pseudotime) of key prioritized TFs in the GC Branch.

**Extended Figure 5:**
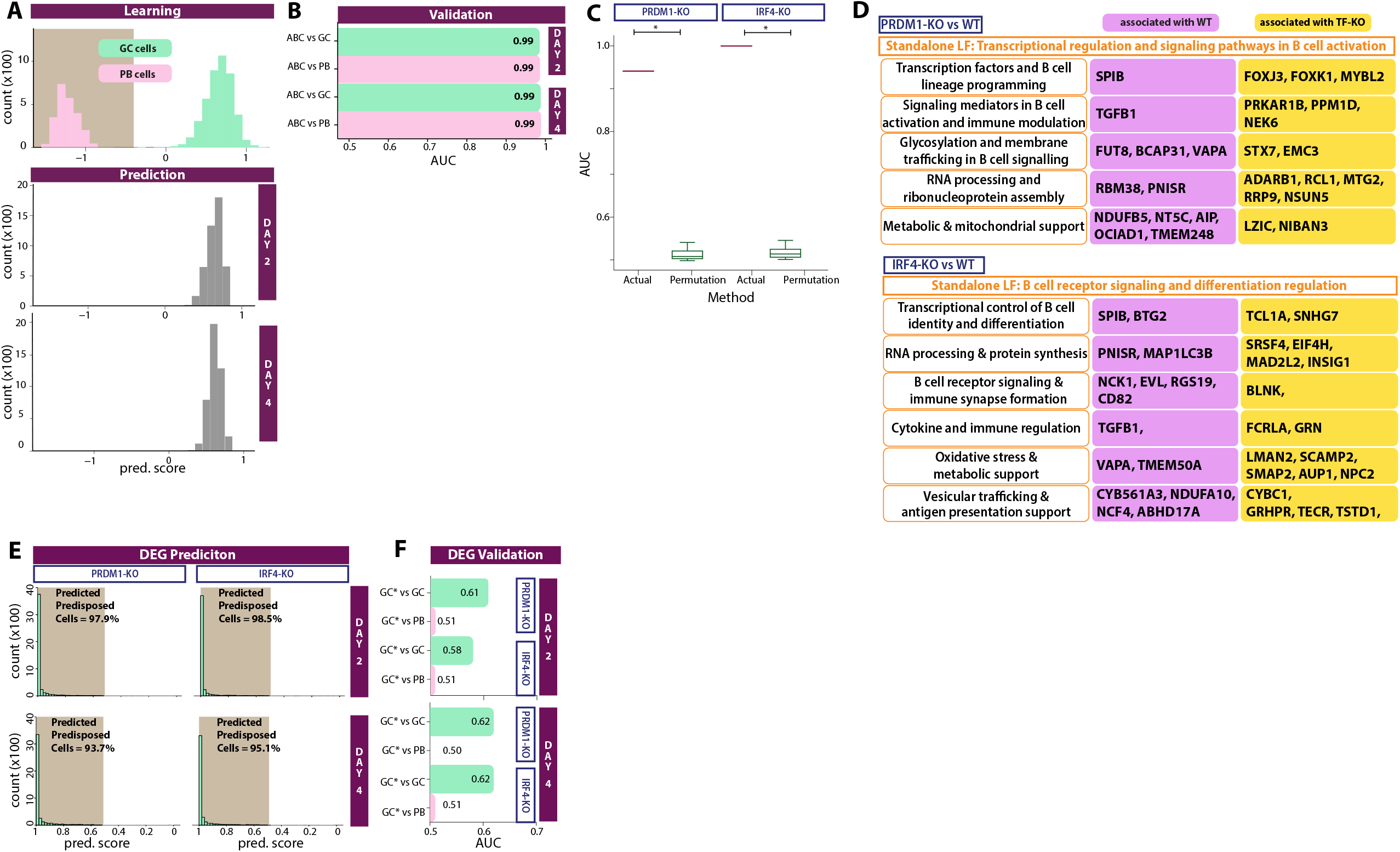
Additional characterization of perturbation-derived LFs that reveal transcriptional predisposition to alternative cell fates. **A**. Projection of LFs learned from the bifurcated GC and PB states onto progenitor populations i.e., unperturbed ABCs at day 2 and day 4. Top panels visualize distribution of LF scores for bifurcated states. Projecting the scores on ABCs at Day 2 and Day 4 (bottom panels) highlights a subset of ABCs (green portion) exhibiting early GC-like fate predisposition at the transcriptional level. **B**. Performance of SLIDE model from A in discriminating predicted GC predisposed cell within the progenitor population against bifurcated GC or PB cell states. **C**. Performance of SLIDE models built to compare PRDM1-KO vs control cells and IRF4-KO and control cells (2 separate models). Model performance evaluated across replicates of k-fold cross-validation and significance evaluated using permutation testing. **D**. Functional annotation of the LFs distinguishing PRDM1-KO and control cells as well as IRF4-KO and control cells. Yellow and pink indicate higher expression in IKZF1-KO/IRF4-KO and control, respectively. **E**. Projection of biomarkers (using DEGs) learned from the PRDM1-KO and IRF4-KO experiments onto progenitor populations i.e., unperturbed ABCs at day 2 and day 4. Projecting the scores on ABCs at Day 2 and Day 4 (bottom panels) highlights a subset of ABCs (green portion) exhibiting early GC-like fate predisposition at the transcriptional level. **F**. Performance of DEGs in discriminating predicted GC predisposed cell within the progenitor population (ABCs at Days 2 and 4) against bifurcated GC or PB cell states.

**Extended Figure 6:**
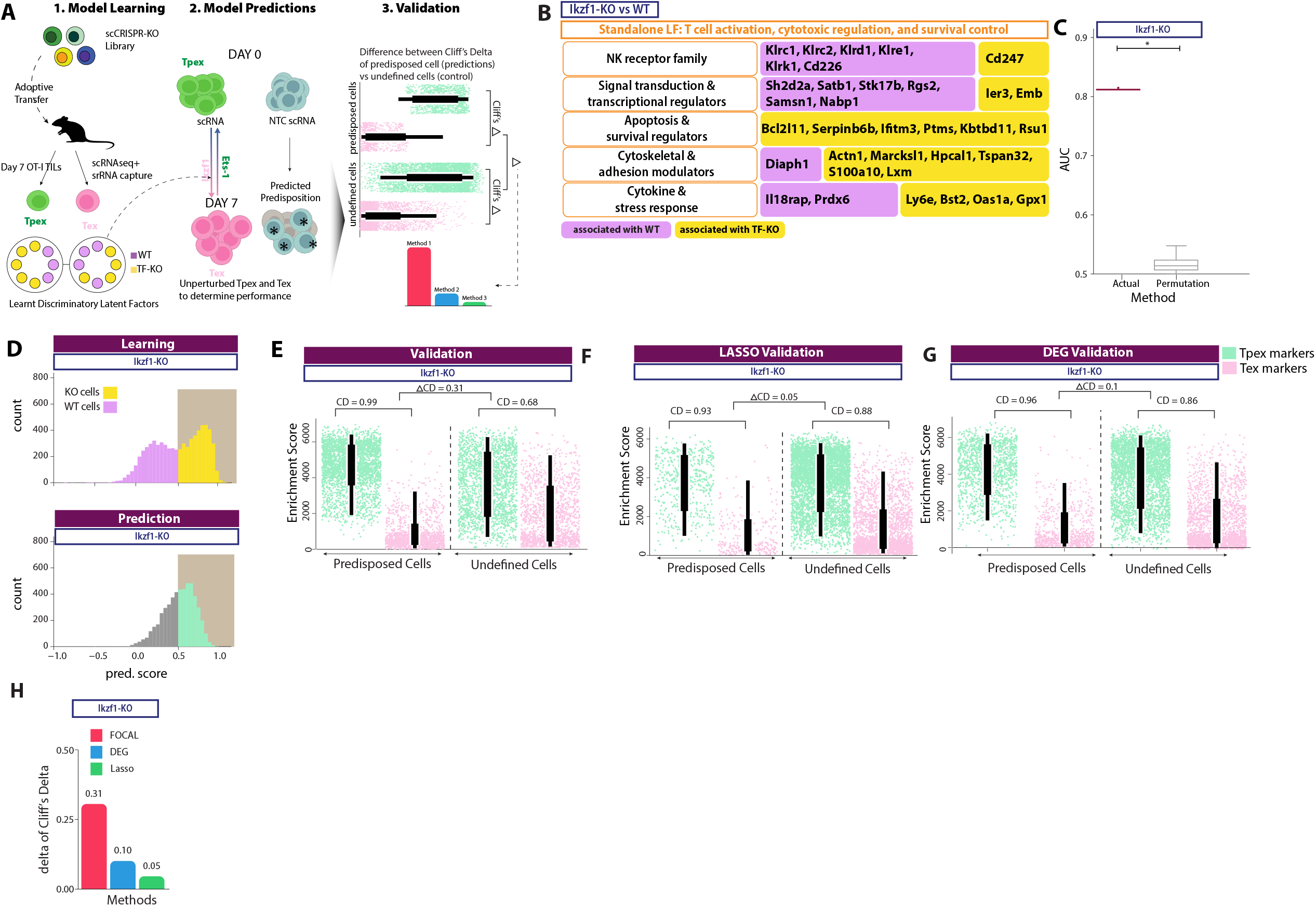
Perturbation-derived latent factors reveal transcriptional predisposition to CD8 T cell exhaustion fates. **A**. LFs learned from bifurcated states in the upper panel and from TF knockout versus unperturbed control cells in the bottom panel are projected onto early ABCs at Day 2 and Day 4 using the multi-ome dataset with the aim to determine whether these programs can predict cell fate in early un-bifurcated cells. Validation is performed by comparing the fraction of early state cells and assessing resembling of their transcriptional profiles to bifurcated cells. **B**. Functional annotation of the LFs distinguishing IKZF1-KO and control cells. Yellow and pink indicate higher expression in IKZF1-KO and control, respectively. **C**. Performance of SLIDE model built to compare IKZF1-KO vs control cells. Model performance evaluated across replicates of k-fold cross-validation and significance evaluated using permutation testing. **D**. Projection of LFs learned from the IKZF1-KO experiment to predict cellular predisposition. Top panels visualize distribution of LF scores for IKZF1-KO cells (yellow) vs unperturbed (purple) cells. Projecting the scores on unperturbed cells highlights a subset of cells exhibiting Tpex-like fate predisposition at the transcriptional level. **E**. ssGSEA enrichment scores reflecting transcriptional predisposition of cells predicted as Tpex-predisposed vs those not predicted as Tpex predisposed by FOCAL. **F**. ssGSEA enrichment scores reflecting transcriptional predisposition of cells predicted as Tpex-predisposed vs those not predicted as Tpex predisposed by a LASSO-based model. **G**. ssGSEA enrichment scores reflecting transcriptional predisposition of cells predicted as Tpex-predisposed vs those not predicted as Tpex predisposed by DE analyses. **H**. Comparison of effect size differences (predicted as predisposed vs not predicted as predisposed) across the 3 methods (FOCAL, LASSO and DE analyses)

## Notes

### Competing Interest Statement

The authors have declared no competing interest.

