## Supplementary material for "Interpretable machine learning coupled to gene regulatory networks uncovers subcircuits underlying cell fate decisions": Combined supplementary notes and figures

### Supplementary Information

#### Supplementary Note 1. Annotating the latent factors using an LLM-based gene-set tool

GSAI was applied to annotate LFs with biological priors. The gene set query to GSAI is provided in Supplementary File S1, with model selection “ChatGPT 5.1”, and prompt “Biological Processes Analysis”.

#### Supplementary Note 2. Construction of the per-window TF-binding prior mask

Each overlapping pseudotime window (Methods, “Pseudo-bulking across trajectory to construct overlapping windows”) is treated as an independent pseudobulk from which one transient-context-specific network is inferred. The binary TF → target prior mask that constrains the edges the network model may fit in that window was constructed in one of two ways, depending on the data available.

*Joint multi-omic dataset.* For each window the RNA cells and ATAC barcodes assigned to that window were selected and read-count QC was reapplied at the thresholds used for the full dataset (Methods). The window’s ATAC fragments were merged and peaks called with MACS2, keeping at most the $5 \times 10^5$ highest-scoring peaks. Transcription-factor (TF) footprints were called inside those peaks with Wellington (pyDNase), supplying the ENCODE blacklist with a maximum of $10^5$ footprints, and the footprints were scanned with HOMER against a CisBP and JASPAR motif collection converted to HOMER format as $\log_2(\text{PWM}/0.25)$ log-odds columns with a per-motif detection threshold. Motif hits were assigned to genes by TSS distance and peak2gene linking, giving a TF-binding table (TF, genomic interval, footprint score) per window together with a binary TF → target prior mask retaining the 20 strongest targets per factor. This mask constrains which edges the network model may fit, so every dynamic edge is supported by an accessible, footprinted motif occurrence in the window in which it appears.

*scRNA alone dataset.* When raw BAM sequence or fragment files from a matched scATAC-seq profile are not present, we utilized CellOracle’s murine and human ATAC-seq derived base GRNs to obtain the binary TF → target prior mask.

#### Supplementary Note 3. Expression, regulation and beta curves along pseudotime

The three curve families below are resolved onto the pseudotime axis of a branch by the NaN-aware Gaussian-kernel smoothing of window-level quantities described in the Methods (“Smoothed regulatory curves along pseudotime”), alongside the force curves defined there.

*Expression curves.* The smoothed $\log_2$ CPM of every gene, $x_g(t)$; the subset restricted to transcription factors supplies the regulator abundance $x_{\text{TF}}(t)$ used by the force definition (Methods). Target-gene responses are shown either as these curves directly or as their pseudotime gradient $dx_g/dt$, which makes the moment of induction or repression of a gene, rather than its absolute level, the quantity being compared across genes.

*Regulation curves.* At each sampled point the smoothed network is binarized by retaining the $k = s \cdot n_{\text{TF}} \cdot n_{\text{target}}$ strongest edges by absolute weight, where $s$ is the network sparsity ($s = 0.01$ throughout), and each TF’s out-degree $d_{\text{TF}}(t)$ in that binarized network is recorded as $\log_2(d_{\text{TF}}(t) + 1)$. This is the number of genes a TF regulates at that point in pseudotime. A weighted variant, in which the out-degree is summed over the weighted rather than the binarized network, is available through the same interface.

*Beta curves.* For a queried set of TF–target links, the signed smoothed edge weight $\beta_{\text{TF} \rightarrow \text{target}}(t)$. Unless stated otherwise the normalized total-effect network ($w_{\text{in}}$) is used; the direct-effect networks ($w_{\text{n}}$, $w$) are available through the same interface. Only the queried sub-network is smoothed.

##### **Supplementary Note 4. Chromatin binding inference across pseudotime trajectories**

TF binding was quantified directly from the per-window chromatin footprints produced during dynamic-network inference. For each window, the inferred TF-to-open-chromatin region (OCR) binding table was read and, for every TF, two per-window quantities were computed: a *binding score*, the mean footprint score of that TF's bound OCRs (averaged within chromosome, then across chromosomes, so that chromosomes with many peaks do not dominate), and a *binding count*, the mean number of bound OCRs per chromosome computed the same way. Windows in which a TF has no bound region contribute a missing score and a count of zero. TFs are either supplied explicitly or discovered as the union of all factors observed across windows.

Each series is then ordered along a branch by that branch's window ordering, mapped onto the branch's pseudotime axis via `AlignTimeScales`, and smoothed with a one-dimensional Gaussian filter ($\sigma = 2$ windows). Because the PB and GC branches are aligned separately, each branch's windows are mapped through its own pseudotime frame. When binding is compared across branches, each TF's curve is optionally min–max scaled using the global minimum and maximum across both branches together, so the two lineages remain on a shared reference scale. These curves give an orthogonal, expression-independent readout of when a factor engages chromatin, and are compared against the force waves of the same factors. The two binding metrics are also compared against each other per TF, with the binding-score and OCR-count curves min–max scaled onto a common axis, to distinguish a genuine strengthening of footprints from a simple increase in the number of accessible regions bound. For comparisons between categories of factors, each lineage's smoothed series is additionally averaged within bins delimited by chosen episode-partition pseudotimes, and each category's per-bin means are compared against a seeded, size-matched random control drawn from factors with a defined binding score in the first bin.

##### **Supplementary Note 5. Optimal-transport couplings and their push-forward and pull-back distributions for ranking transcription factors**

FocalFire prioritizes transcription factors (TFs) from its dynamic gene regulatory networks. As an independent ranking that uses no network, pseudotime or force model, we computed optimal-transport (OT) couplings between the consecutive sampling windows of the B-cell culture with `moscot` [cite]. A coupling assigns every cell of one window a distribution over the cells of the next, and from it two per-cell quantities are derived: the *pull-back* of a cell state, which weights the cells of the earlier window by how much of that state they give rise to, and the *push-forward* of a cell state, which weights the cells of the later window by how much of that state they descend from. Correlating TF expression with each of these distributions ranks the TFs in two complementary ways. This note gives only the OT background needed to define the coupling, defines the push-forward and pull-back distributions, and states how the TF rankings were computed from them; for the method itself we refer to `moscot`.

###### ***Optimal-transport coupling between consecutive time points***

Consider two consecutive time points $t_k < t_{k+1}$ with $n$ cells profiled at $t_k$ and $m$ cells at $t_{k+1}$. moscot treats the two sets of cells as discrete probability measures with marginal weights $a \in \mathbb{R}^n$ and $b \in \mathbb{R}^m$; we used uniform marginals, $a_i = 1/n$ and $b_j = 1/m$, so that every cell carries the same mass. Given a cost $C_{ij}$ for moving cell $i$ to cell $j$, moscot solves the entropically regularized OT problem

$$\begin{aligned} P^* &= \arg \min_{P \geq 0, P\mathbf{1}=a, P^\top \mathbf{1}=b} \sum_{i,j} P_{ij} C_{ij} - \varepsilon H(P), \quad H(P) \\ &= - \sum_{i,j} P_{ij} (\log P_{ij} - 1), \end{aligned} \quad (\text{S1})$$

where $\varepsilon > 0$ sets how diffuse the solution is. The minimizer $P^* \in \mathbb{R}^{n \times m}$ is the *coupling*: $P_{ij}^*$ is the probability mass carried from cell $i$ at $t_k$ to cell $j$ at $t_{k+1}$, with rows summing to $a$ and columns to $b$. Dividing the coupling by its marginals gives two conditional matrices,

$$F_{ij} = \frac{P_{ij}^*}{a_i}, \quad B_{ij} = \frac{P_{ij}^*}{b_j}, \quad (\text{S2})$$

where row $i$ of $F$ sums to one and is the distribution of cell $i$ over the cells of $t_{k+1}$ (where its mass goes), and column $j$ of $B$ sums to one and is the distribution of cell $j$ over the cells of $t_k$ (where its mass comes from). All quantities below are built from $F$ and $B$.

**Transport cost.** Rather than the Euclidean distance between two cells in an embedding, which can cut straight across regions that contain no cells, we used moscot’s graph geodesic cost. A 30-nearest-neighbour graph was built on the cells of both time points together, and the cost between two cells was taken as

$$C_{ij} = -4\tau \log(H_\tau)_{ij}, \quad (\text{S3})$$

where $H_\tau$ is the heat kernel of the graph at diffusion time $\tau$ ($\tau = 100$). This approximates the squared distance along the data manifold, so two cells are close only if the graph connects them through intermediate cells. The graph was built on the joint RNA/ATAC latent space learned with MultiVI [cite] from the cells with paired scRNA-seq and scATAC-seq profiles (Methods). Defining the cost on this joint space means that two cells are close only if they are similar in both transcriptome and chromatin accessibility, so the coupling cannot connect cells that share expression but differ in regulatory state.

**Couplings computed.** Within each sampling window, cells were restricted to the cell states populating that stage of the culture (day 0–2: earlyActB, ActB-1, ActB-2; day 3–4: ActB-2, ActB-3, ActB-4, earlyPB; day 5–6: GC-1, GC-2, PB-2), leaving 28,243 cells ($n = 9,850, 9,282$ and $9,111$ per window), and the windows were encoded as $t_1 = 1.5$, $t_2 = 3.5$ and $t_3 = 5.5$. The problems $t_1 \rightarrow t_2$ and $t_2 \rightarrow t_3$ were solved with $\varepsilon = 10^{-3}$. We write $P^{(k)}$, $F^{(k)}$ and $B^{(k)}$ for the coupling and conditional matrices of the pair $(t_k, t_{k+1})$; the TF rankings use the second coupling, $k = 2$.

#### **Push-forward and pull-back distributions**

The coupling is a linear map between weight vectors on the two sets of cells. Given weights $u \in \mathbb{R}_{\geq 0}^n$ on the cells at $t_k$, their *push-forward* is the weight vector they induce on the cells at $t_{k+1}$; given weights $v \in \mathbb{R}_{\geq 0}^m$ on the cells at $t_{k+1}$, their *pull-back* is the weight vector they induce on the cells at $t_k$:

$$\begin{aligned} (\text{push}_k(u))_j &= \sum_i F_{ij}^{(k)} u_i = \sum_i \frac{P_{ij}^{(k)}}{a_i} u_i, & (\text{pull}_k(v))_i &= \sum_j B_{ij}^{(k)} v_j \\ &= \sum_j \frac{P_{ij}^{(k)}}{b_j} v_j. \end{aligned} \quad (\text{S4})$$

Each earlier cell contributes its own forward distribution (row $i$ of $F$) with weight $u_i$, and each later cell contributes its own backward distribution (column $j$ of $B$) with weight $v_j$. For a cell state $S$ with indicator vector $\mathbf{1}_S$ and $|S|$ cells, $\text{push}_k(\mathbf{1}_S/|S|)$ is a probability distribution over the later cells that gives each cell $j$ its probability of having come from $S$, and $\text{pull}_k(\mathbf{1}_S/|S|)$ is a probability distribution over the earlier cells that gives each cell $i$ its probability of contributing to $S$. Using the unnormalized indicator $\mathbf{1}_S$ scales these vectors by $|S|$ without changing the ordering of cells; with uniform marginals both are constant multiples of the raw coupling sums $\sum_{i \in S} P_{ij}^{(k)}$ and $\sum_{j \in S} P_{ij}^{(k)}$.

#### **Ranking transcription factors with the two distributions**

Let $c_{gi}$ be the raw UMI count of gene $g$ in cell $i$. Expression was normalized to $10^4$ counts per cell and log-transformed,

$$E_{gi} = \log \left( 1 + 10^4 \frac{c_{gi}}{\sum_{g'} c_{g'i}} \right), \quad (\text{S5})$$

with $g$ ranging over the TFs of moscot's built-in human TF annotation present in the expression matrix. Given a per-cell score $s_i$ defined on a set of cells $\mathcal{C}$, every TF was scored by its Pearson correlation with $s$ over $\mathcal{C}$,

$$r_g = \frac{\sum_{i \in \mathcal{C}} (E_{gi} - \bar{E}_g) (s_i - \bar{s})}{\sqrt{\sum_{i \in \mathcal{C}} (E_{gi} - \bar{E}_g)^2} \sqrt{\sum_{i \in \mathcal{C}} (s_i - \bar{s})^2}}, \quad (\text{S6})$$

where $\bar{E}_g$ and $\bar{s}$ are means over $\mathcal{C}$. A two-sided $p$-value $p_g$ was obtained from the Fisher $z$-transform of $r_g$ and corrected across TFs by the Benjamini–Hochberg procedure to give $q_g$. TFs were ranked by $r_g$ and called significant when

$$q_g < 0.05 \quad \text{and} \quad |r_g| > 0.1. \quad (\text{S7})$$

The two rankings differ only in the score $s$ and the cell set $\mathcal{C}$. Both were computed for the source state $A = \text{ActB-4}$ at $t_2$ (day 3–4) and the target state $B = \text{GC-2}$ at $t_3$ (day 5–6).

**Pull-back ranking.** The target state is pulled back through the second coupling to the day 3–4 cells, and every day 3–4 cell $i$ receives the score

$$s_i^{\leftarrow} = \mathbf{1}_A(i) + (\text{pull}_2(\mathbf{1}_B))_i = \mathbf{1}_A(i) + \sum_{j \in B} B_{ij}^{(2)}, \quad \mathcal{C} = \{\text{cells at } t_2\}, \quad (\text{S8})$$

whose first term marks membership of the source state and whose second term is the mass of $B$ that cell $i$ contributes. TFs with positive $r_g$ are expressed in the day 3–4 cells that feed $B$, and TFs with negative $r_g$ in the day 3–4 cells that do not; this ranking yielded 38 significant TFs.

**Push-forward ranking.** The uniform distribution over the source state is pushed forward through the second coupling to the day 5–6 cells, so that every day 5–6 cell $j$ receives its probability of having come from $A$,

$$s_j^{\rightarrow} = (\text{push}_2(\mathbf{1}_A/|A|))_j = \frac{1}{|A|} \sum_{i \in A} F_{ij}^{(2)}, \quad \mathcal{C} = B, \quad (\text{S9})$$

and the correlation is taken over the cells of $B$ only. TFs with positive $r_g$ are expressed in the $B$ cells that came from $A$, and TFs with negative $r_g$ in the $B$ cells that came from elsewhere; this ranking yielded 8 significant TFs.

#### Supplementary Note 6. Gene set enrichment analysis using ESCAPE

Single-cell gene set enrichment was performed using ESCAPE (Enrichment of Single Cell Analysis by Pathway Expression) (ref) to assess whether predicted KO-like subsets exhibited transcriptional programs consistent with the expected lineage bias. Enrichment scores were calculated using predefined marker sets for progenitor-like exhausted T cells (Tpex: *Tcf7*, *Slamf6*, *Myb*, *Sell*, and *Bach2*) and terminally exhausted T cells (Tex: *Havcr2*, *Pdcd1*, *Entpd1*, *Cd38*, *Cd244a*, *Cxcr6*, and *Ifng*).

### **SUPPLEMENTARY FIGURE LEGENDS**

#### **Supplementary Fig 1: Imputation of sc-RNA seq data from multi-omic B cell dataset**

- **A.** Pre-imputed expression of lineage-defining TFs across the B cell UMAP.
- **B.** PCA embedding of the B cell multiome data colored by sub-cell state.
- **C.** Diffusion map embedding colored and labeled by sub-cell state. Cells are ordered from naive and early activated states through the activated compartment toward the GC and PB endpoints using first two diffusion map eigenvectors (X and Y axes).
- **D.** Post filtering UMAP embedding with sub-cell state annotations.
- **E.** Expression of the same TFs as in A after MAGIC imputation, plotted on UMAP with the same color scale.

#### **Supplementary Fig 2: Latent Factor score and TF expression along the B cell branching trajectory.**

- **A.** Interacting latent factor Z3 score across the STREAM branching trajectory. S1 marks the trajectory origin and S2 (PB) and S3 (GC) mark the two terminal branches; the arrow indicates the direction of increasing pseudotime.
- **B–H.** Expression of key lineage-defining TFs along the same branching trajectory.

#### **Supplementary Fig 3: Trajectory reconstruction of the Tpex-to-TeX transition.**

##### **A. Cell state composition across binned pseudotime windows.**

Average cell counts per sliding window (n = 500 cells per overlapping window) along the Tpex to Tex trajectory, with Tpex (dark blue) and Tex (pink) from Bin 1 to Bin 4.

- **B.** Population dynamics across the Tpex-to-TeX branch pseudotime.
- **C.** STREAM elastic principal graph learned in LLE space for the linear trajectory.
- **D.** Cells colored by state (dark blue: Tpex, pink: Tex) with the fitted principal curve running from the S0 origin of Tpex to the S1 terminus of Tex.

#### **Supplementary Fig 4: Expression of canonical effector and exhaustion genes along the Tpex-to-TeX pseudotime.**

##### **A–F. Time-resolved expression of marker genes across the T cell trajectory.**

Per-cell expression of TFs plotted on the STREAM subway map from the Tpex (S0) origin to the Tex (S1) terminus; the arrow indicates the direction of increasing pseudotime.

#### **Supplementary Fig 5: Compositional data for TF Force derivation**

- **A.** Heatmap of indirect effect beta-values of enriched links from FOCAL-GIFs across pseudotime for the PB branch, with red indicating positive and blue negative indirect effects.
- **B.** Smoothed expression (CPM) across pseudotime for the TFs driving the edges in A.
- **C.** As in A, indirect effect beta-values for prioritized TF-target edges in the GC branch.
- **D.** Smoothed expression (CPM) across pseudotime for the TFs driving the edges in C.

#### **Supplementary Fig 6: Episodic GRN edge force distributions along the GC B cell trajectory.**

**A–H.** Histograms of TF force averaged across the temporal duration of an episode (x-axis), of all edges in the episodic GRN. Panels display inference for each of the eight consecutive episodes spanning the ActB-1 to late GC transition, separated into positive (red) and negative (blue) forces.

**Supplementary Fig 7: Per-episode GRN edge force distributions along the T cell exhaustion trajectory.**

**A–F.** Histograms of the average force across the temporal duration of an episode (x-axis) of all edges in the episodic GRN inferred for each of the six consecutive episodes spanning the Tpex to Tex transition, separated into positive (red) and negative (blue) forces.

**Supplementary Fig 8: TF binding scores and chromatin accessibility across pseudotime in B cells**

**A.** Chromatin dynamics plotted along pseudotime for each TF, with solid lines denoting the binding score and dashed lines the Open Chromatin Regions (OCRs) on the same relative scale.

**B–C.** TF expression and downstream number of gene dynamics across pseudotime for key-enriched TFs.

**Supplementary Fig 9: SCENIC+ eRegulon benchmarking in GC and PB B cells.**

UMAP displays scRNA-seq cell embeddings colored by state (blue: GC, orange: PB). Dot plot summarizes eRegulon specificity across the two states, separated into activator and repressor modules, where color denotes the gene-based regulon AUC of the two states, and dot size the corresponding region-based enrichment. Per-cell AUC scores for eRegulons of each TF are projected onto the same embedding, with the number of target genes per eRegulon given in parentheses.

**Supplementary Fig 10: SCENIC+ eRegulon benchmarking in Tex-KLR and Tex-term CD8<sup>+</sup> T cells.** See the description of Supplementary Fig 9, here applied to CD8<sup>+</sup> T cells at the exhaustion bifurcation.

**Supplementary Fig 11: Dictys benchmark for prioritizations of TFs along the bifurcating B-cell trajectory**

**A.** Smoothed TF expression curves (left) and TF regulatory out-degree curves (right) across pseudotime for the PB branch.

**B.** As in A, for the GC branch, showing the corresponding switching and transient TFs.

**Supplementary Fig 12: Episodic TF forces on PRDM1-KO and IRF4-KO latent factors**

**A.** Heatmap of TF force strength exerted by IRF8 and NFATC2 on PRDM1-KO LF genes across the eight episodes spanning the ActB-1 to late GC trajectory. Entries are colored yellow for positive and blue for negative forces, and blank cells indicate episodes in which the edge did not pass the significance threshold. Target genes are grouped by the functional module assigned to the program (cellular trafficking and post-translational modification).

**B.** As in A, IRF8 and NFATC2 act on IRF4-KO LF genes, grouped into protein trafficking and cellular localization, cellular stress response and signaling modules.

**Supplementary Fig 13: A.** GSAI derived categories of phenotypic signatures for latent factor genes associated with ETS1-KO (yellow) or WT (pink) cells.

B. SLIDE model performance after cross-validation over K-data splits for label prediction.

C. FOCAL learning displays the z-score distribution with the split between WT and KO cells, and FOCAL prediction is on the un-perturbed cells using the same z-loadings.

D-F. ESCAPE based scoring of cells to stratify the difference in cell populations as predicted by SLIDE transfer learning, LASSO and DEGs. Differences between enrichment scores are quantified using cliff's delta for predicted predisposed and undefined populations.

G. The difference of cliff's delta between the three approaches is plotted.

H-I. State-specific and Episodic GIFs using transfer learnt latent factors reveal key TFs.

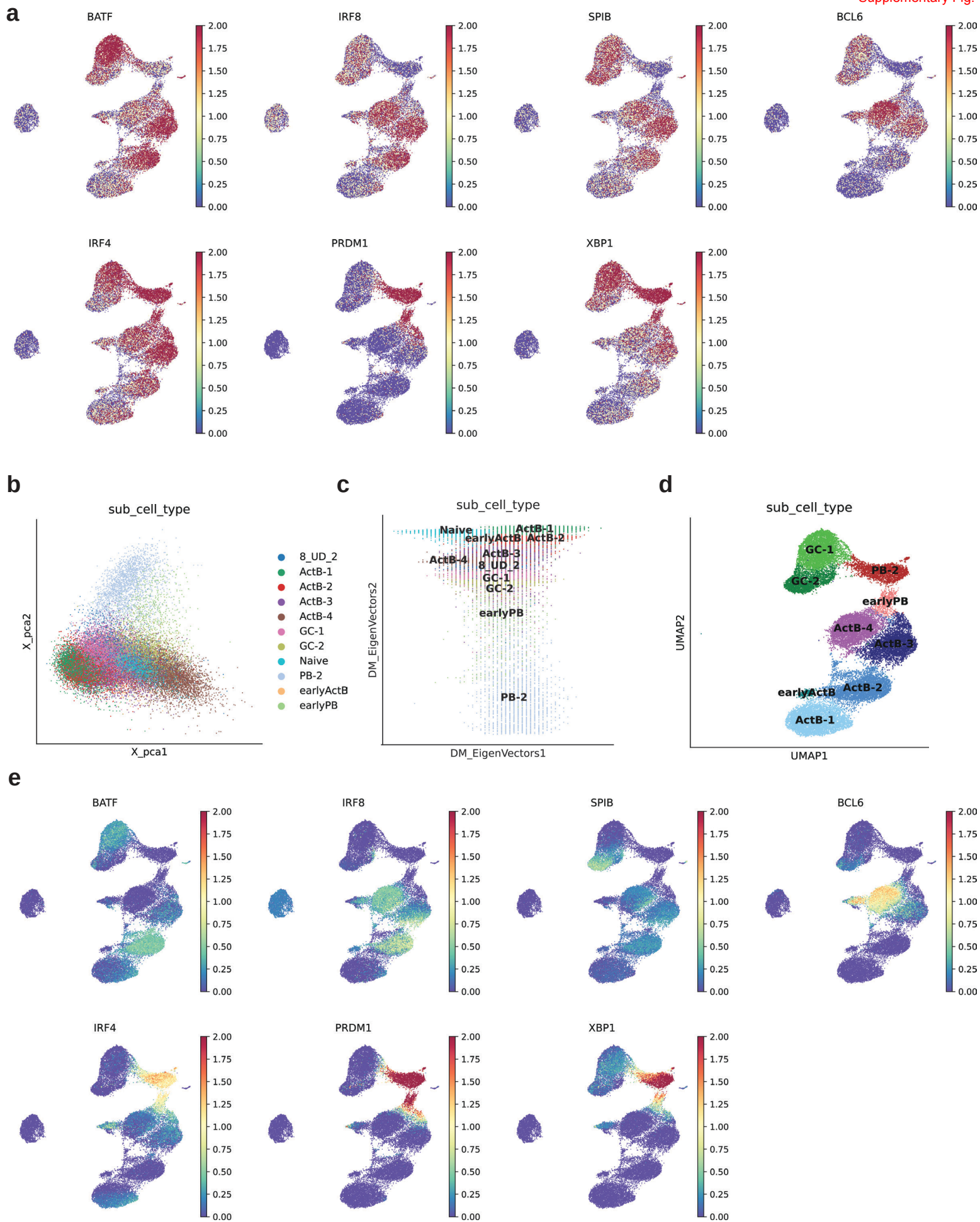

**a**

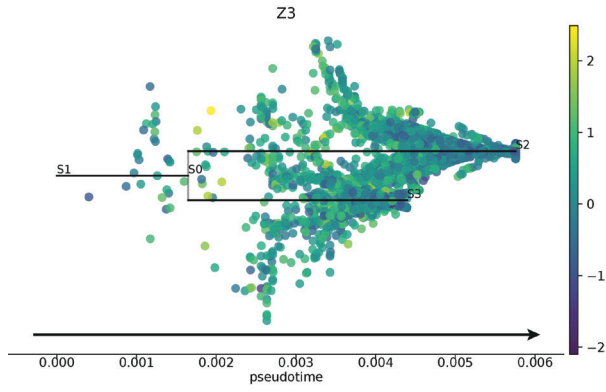

**b**

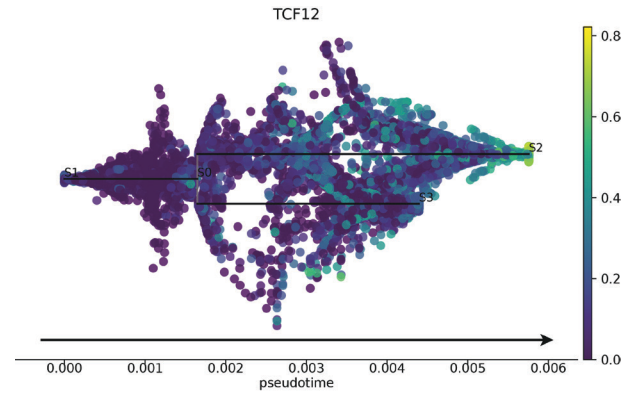

**c**

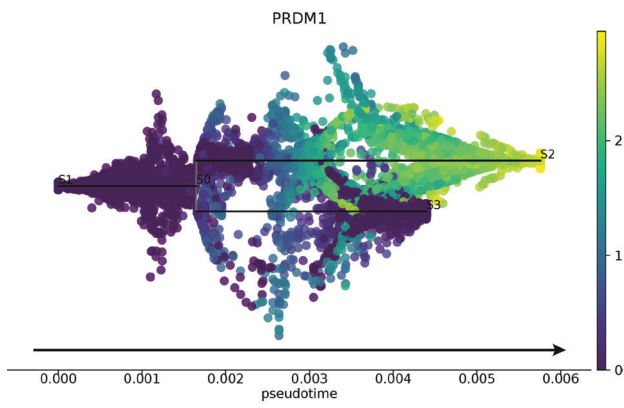

**d**

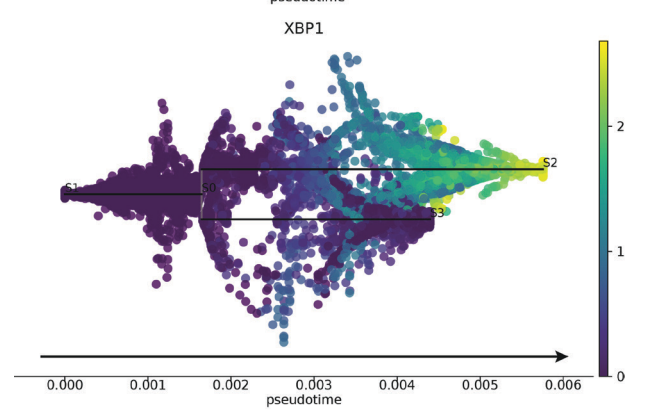

**e**

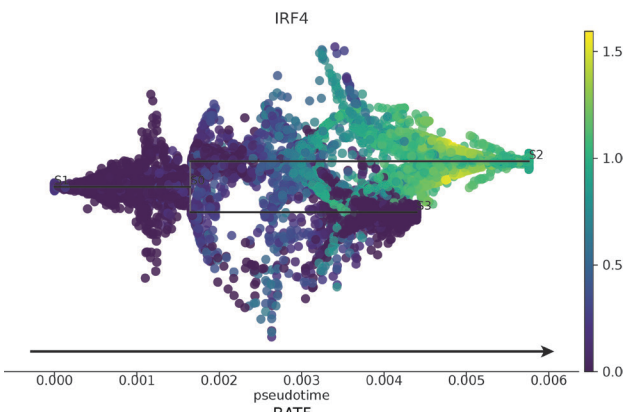

**f**

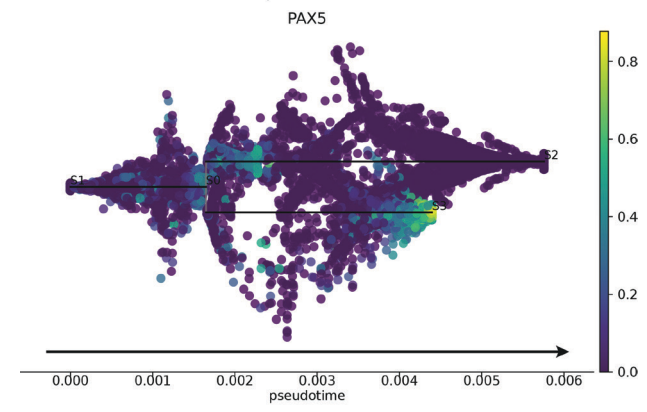

**g**

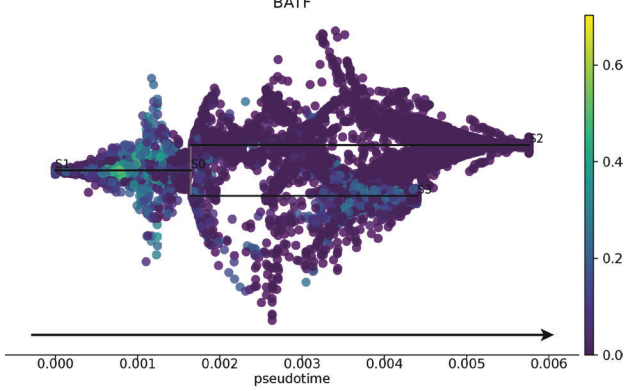

**h**

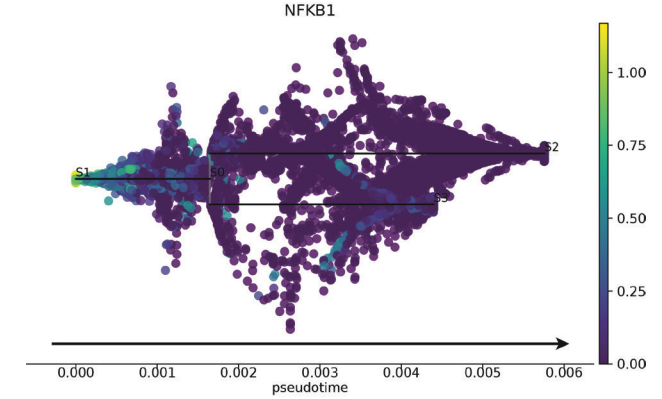

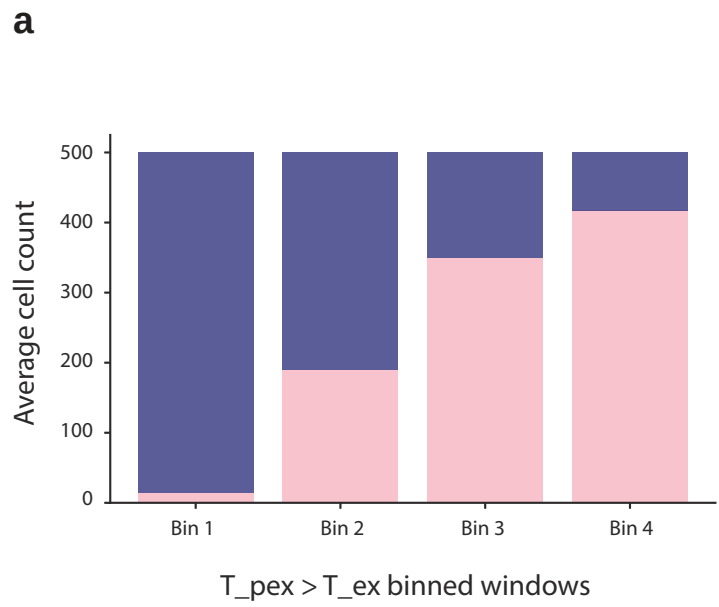

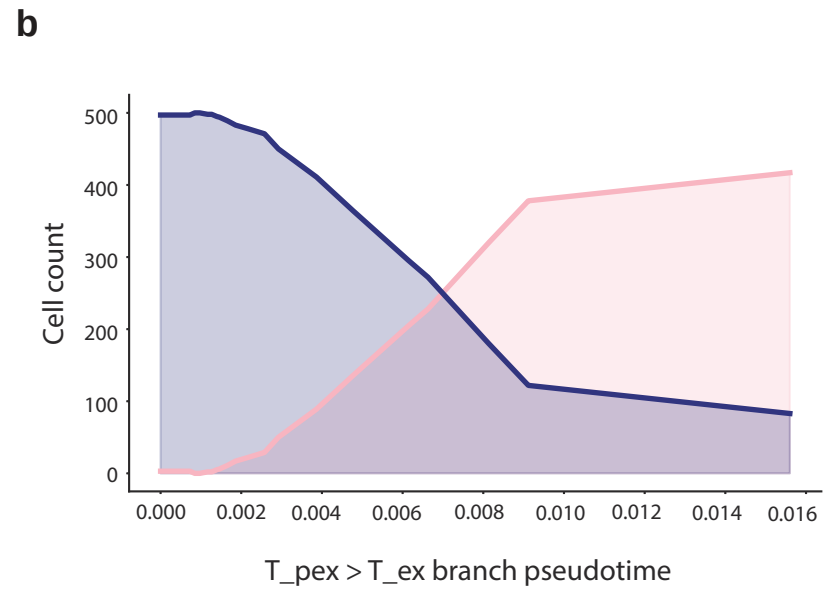

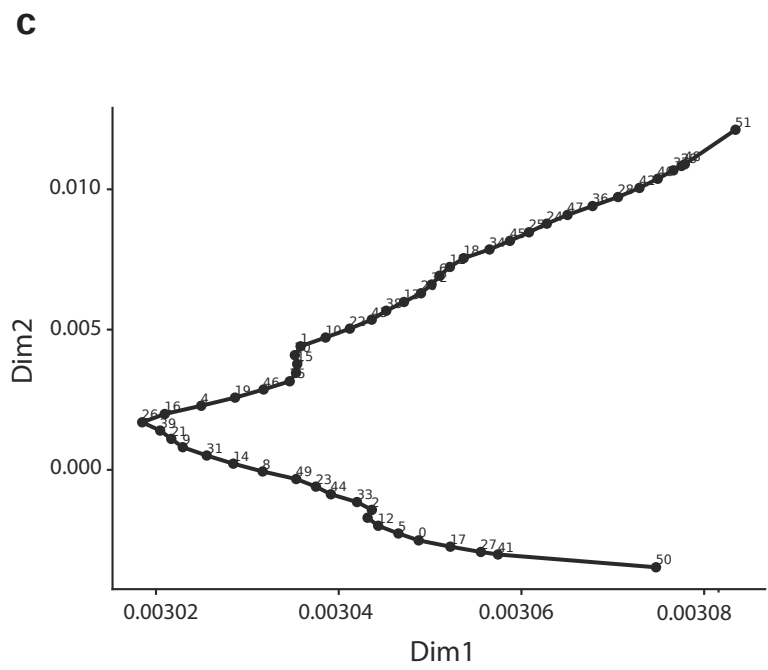

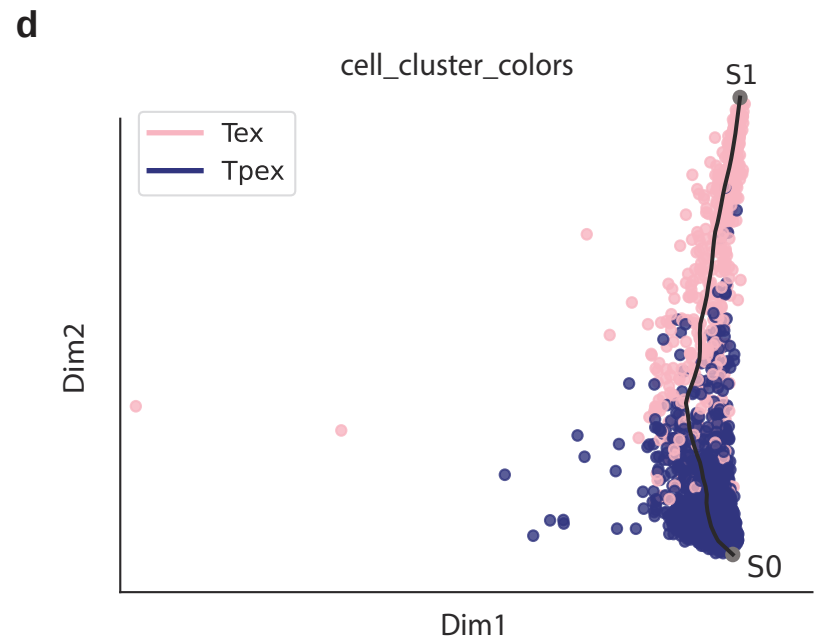

**a**

Gzmb

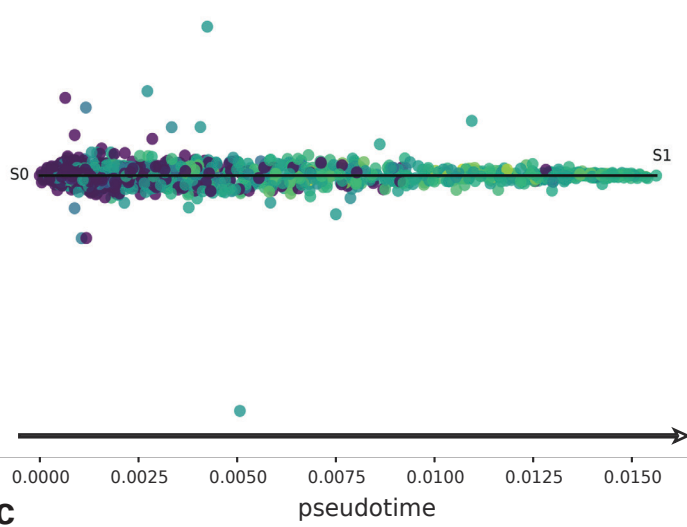

**b**

Gzma

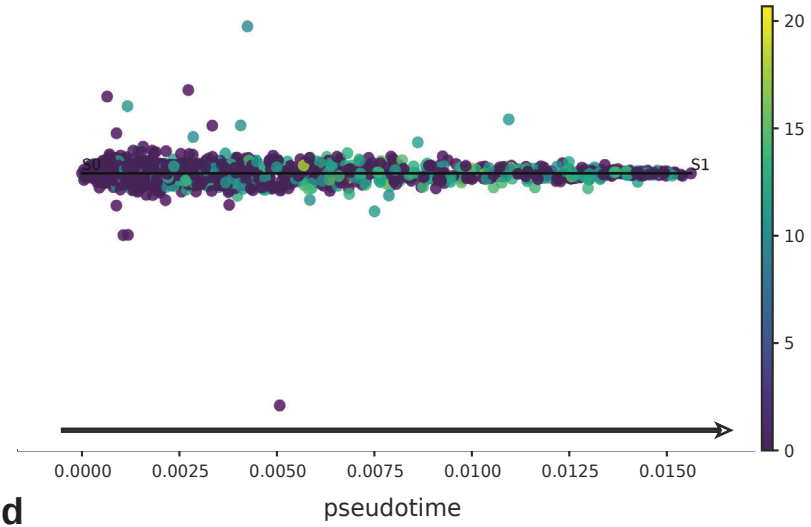

**c**

Klf2

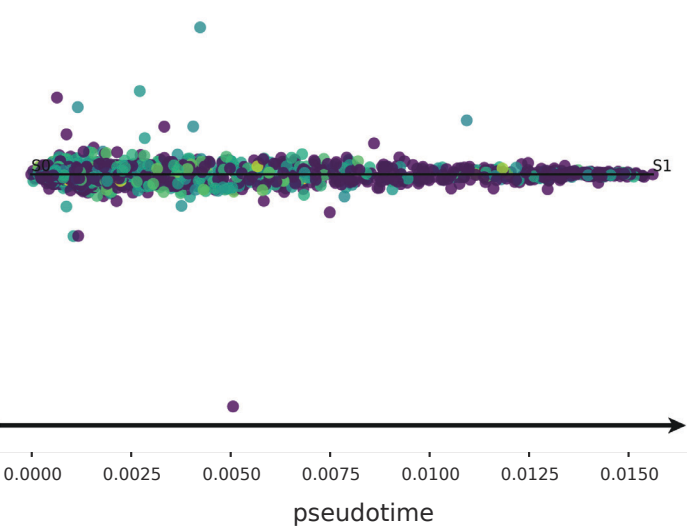

**d**

E2f2

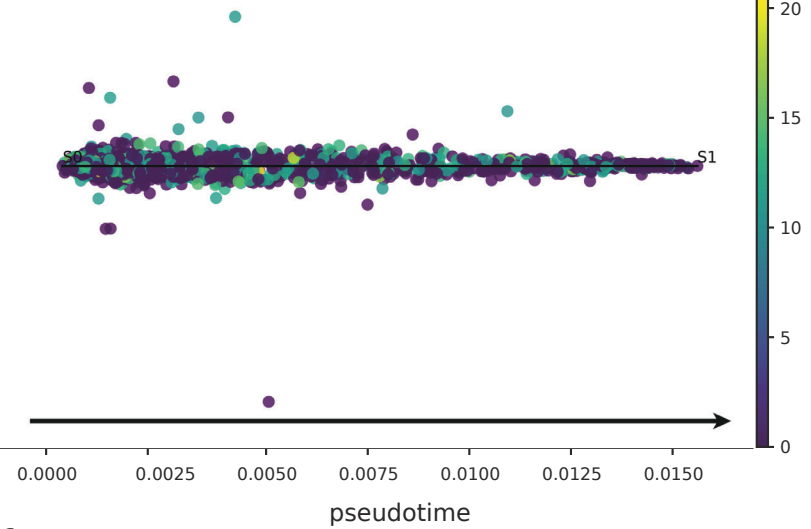

**e**

Tox

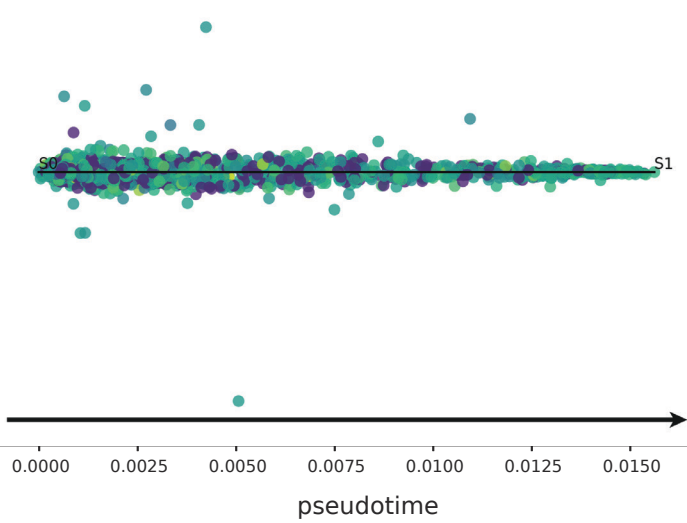

**f**

Foxo1

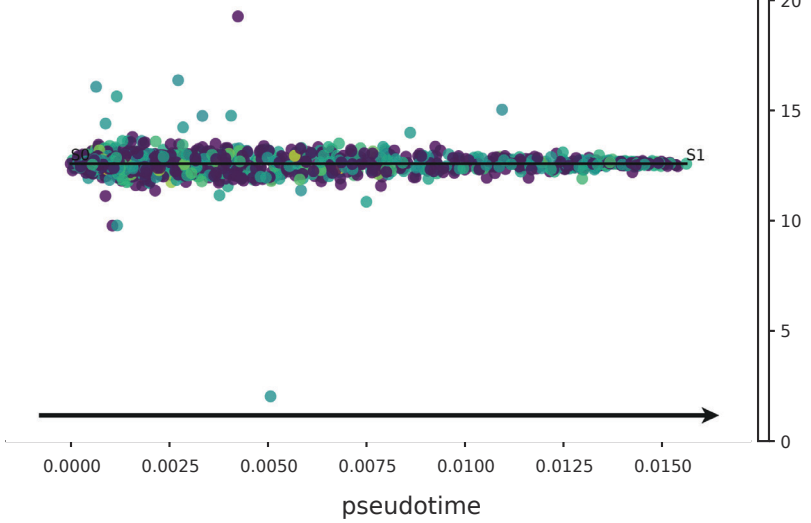

**a**

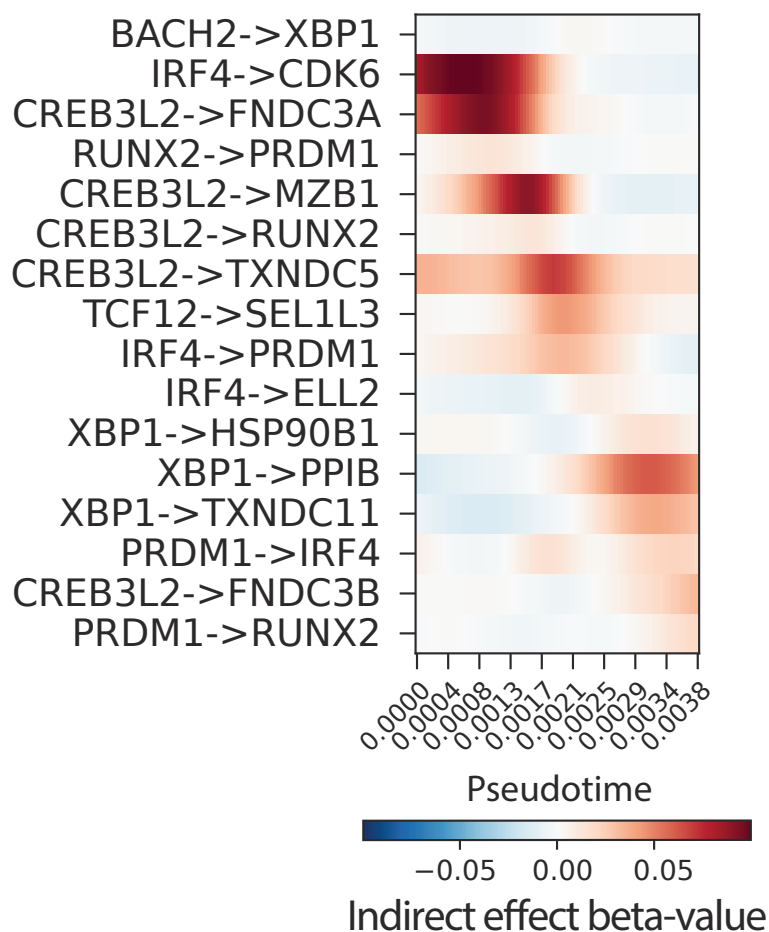

**b**

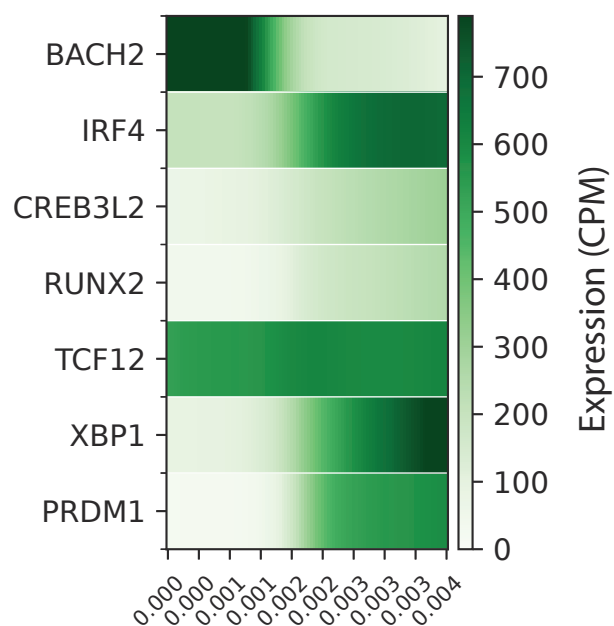

**c**

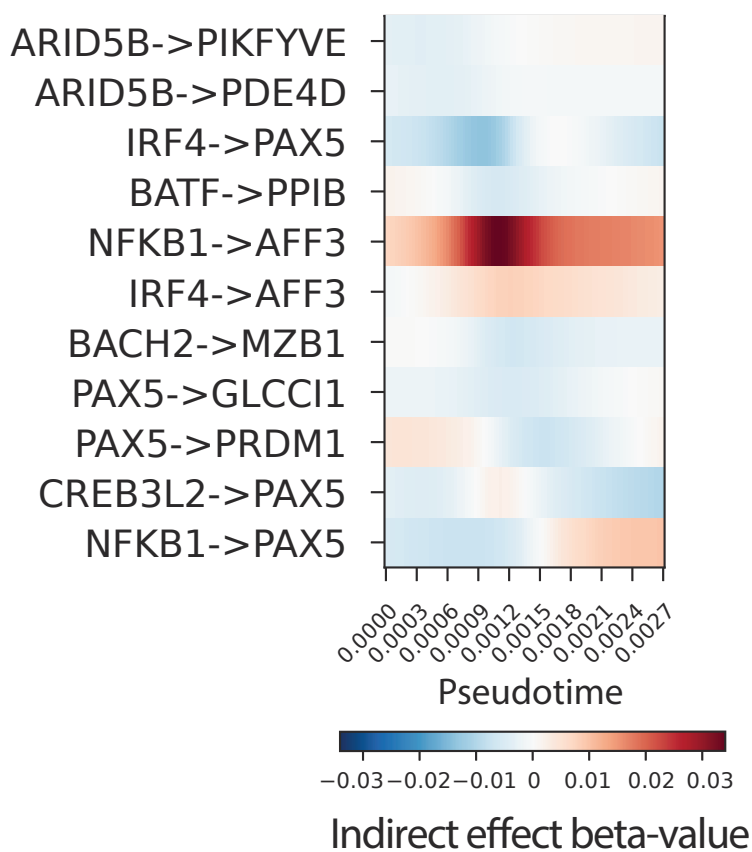

**d**

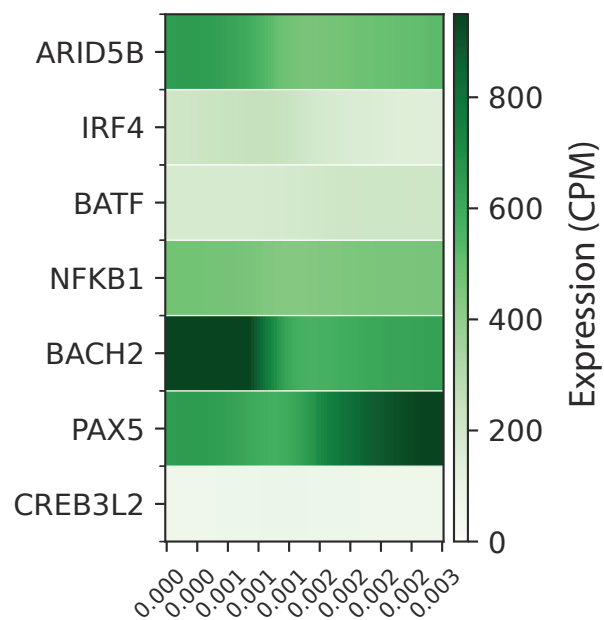

### Episodic GRN edge distributions along trajectory of cellular transitions from Activated Day-1 to late GC B-cell states

**a**

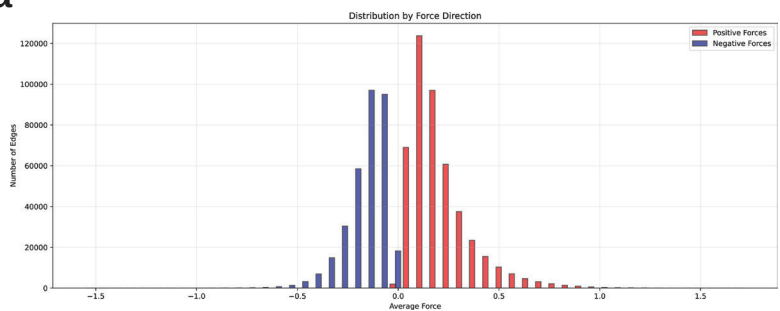

Edge distribution of Episode 1 (ActB-1 > GC)

**b**

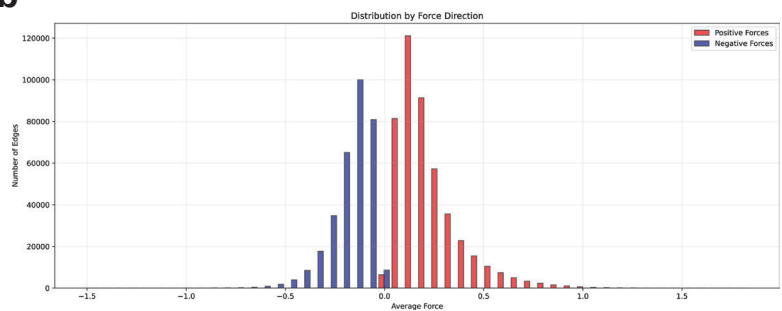

Edge distribution of Episode 2 (ActB-1 > GC)

**c**

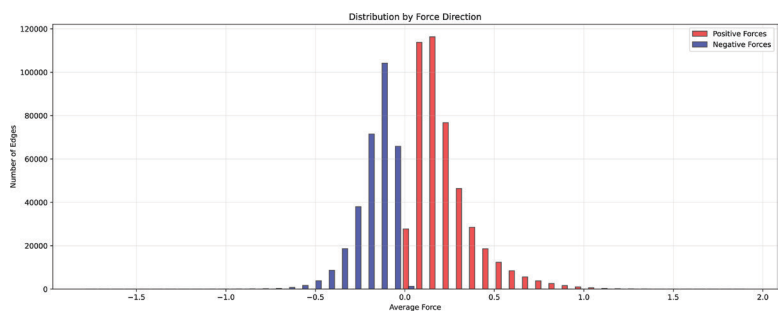

Edge distribution of Episode 3 (ActB-1 > GC)

**d**

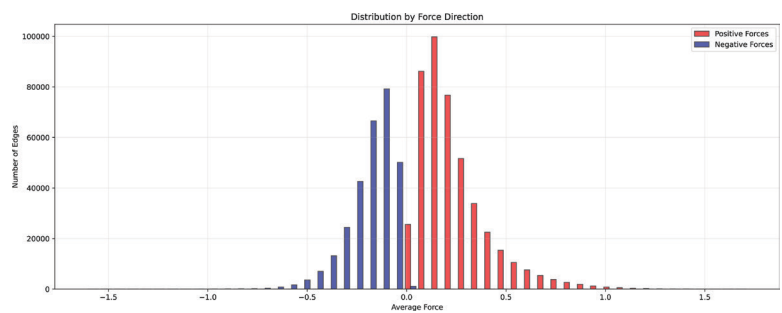

Edge distribution of Episode 4 (ActB-1 > GC)

**e**

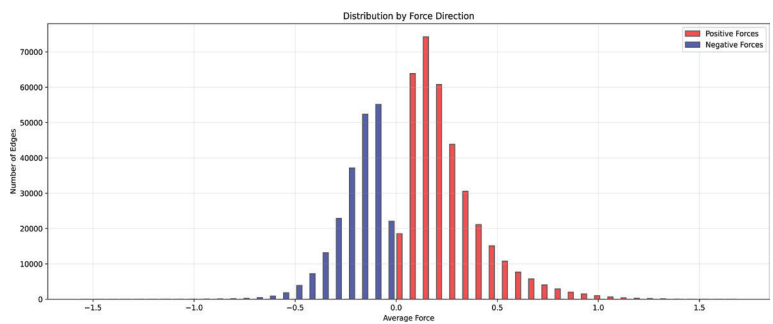

Edge distribution of Episode 5 (ActB-1 > GC)

**f**

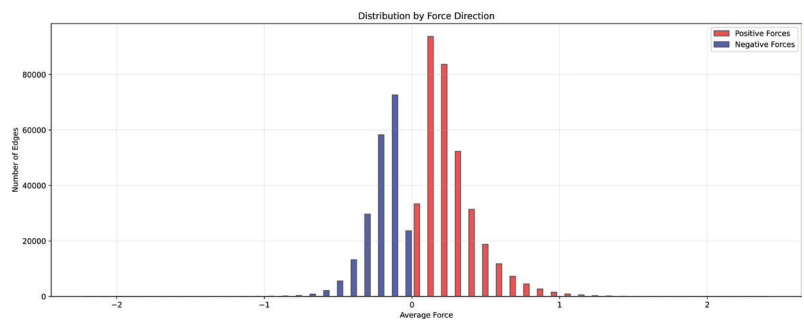

Edge distribution of Episode 6 (ActB-1 > GC)

**g**

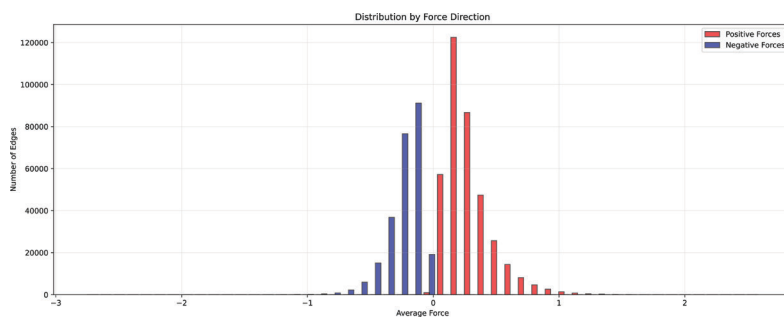

Edge distribution of Episode 7 (ActB-1 > GC)

**h**

Edge distribution of Episode 8 (ActB-1 > GC)

### Per Episode GRN edge distributions along trajectory of cellular transitions from pre-Exhausted to Exhausted T-cell states

**a**

Episode 1

**b**

Episode 2

**c**

Episode 3

**d**

Episode 4

**e**

Episode 5

**f**

Episode 6

### TF OCR counts and Binding score (GC branch)

OCR counts > TF binding score region
TF binding score
TF OCR counts

SCENICplus on B cells

Supplementary Fig. 9

SCENICplus on T cells

direct\_gene\_based\_AUC

direct\_region\_based\_AUC

**a**

**Plasma Blast branch -**

TF expression curves

TF regulatory out-degree curves

**b**

**Germinal Center branch -**

TF expression curves

TF regulatory out-degree curves

a

b
